# Development of a Well-Characterized Cynomolgus Macaque Model of Sudan Virus Disease for Support of Product Development

**DOI:** 10.1101/2022.05.04.490569

**Authors:** Kendra J. Alfson, Yenny Goez-Gazi, Michal Gazi, Ying-Liang Chou, Nancy A. Niemuth, Marc E. Mattix, Hilary Staples, Benjamin Klaffke, Gloria F. Rodriguez, Priscilla Escareno, Carmen Bartley, Anysha Ticer, Elizabeth A. Clemmons, John W. Dutton, Anthony Griffiths, Gabe T. Meister, Daniel C. Sanford, Chris M. Cirimotich, Ricardo Carrion

## Abstract

The primary objective of this study was to characterize the disease course in cynomolgus macaques exposed to Sudan virus (SUDV), to determine if infection in this species is an appropriate model for the evaluation of filovirus countermeasures under the FDA Animal Rule. Sudan virus causes Sudan virus disease (SVD), with an average case fatality rate of approximately 50%, and while research is ongoing, presently there are no approved SUDV vaccines or therapies. Well characterized animal models are crucial for further developing and evaluating countermeasures for SUDV. Twenty (20) cynomolgus macaques were exposed intramuscularly to either SUDV or sterile phosphate buffered saline; 10 SUDV-exposed animals were euthanized on schedule to characterize pathology at defined durations post-exposure and 8 SUDV-exposed animals were not part of the scheduled euthanasia cohort. Survival was assessed, along with clinical observations, body weights, body temperatures, hematology, clinical chemistry, coagulation, viral load (serum and tissues), macroscopic observations, and histopathology. There were statistically significant differences between SUDV-exposed animals and mock-exposed animals for 26 parameters, including telemetry body temperature, clinical chemistry parameters, hematology parameters, activated partial thromboplastin time, serum viremia, and biomarkers that characterize the disease course of SUDV in cynomolgus macaques.

## Introduction

Filoviruses are non-segmented, single-stranded, negative-sense RNA viruses, and some members of the family are known to infect humans and nonhuman primates (NHPs) with severe health consequences [1, 2]. Within the family Filoviridae, there are now five genera: *Ebolavirus*, *Marburgvirus*, *Cuevavirus*, *Striavirus*, and *Thamnovirus*; *Dianlovirus* is proposed as a sixth. There are six species in the *Ebolavirus* genus: *Zaire ebolavirus* (Ebola virus, EBOV), *Sudan ebolavirus* (Sudan virus, SUDV), *Taï Forest ebolavirus* (Taï Forest virus, TAFV), *Reston ebolavirus* (Reston virus, RESTV), *Bundibugyo ebolavirus* (Bundibugyo virus, BDBV), and *Bombali ebolavirus* (Bombali virus, BOMV) [3–5].

EBOV has been a primary target for filovirus research due to high mortality and the large 2014-2016 outbreak in western Africa [6]. However, there are now preventative and therapeutic measures available for EBOV [7–9]. SUDV causes Sudan virus disease (SVD) [5], with an average case fatality rate of approximately 50% [1, 2, 10, 11], and infection with less than one infectious particle is reported to be sufficient to cause disease in NHPs [12]. While research is ongoing, with many candidates in the pipelines, presently there are no approved SUDV vaccines or therapies [13]. Well characterized animal models are crucial for further developing and evaluating countermeasures for SUDV.

Characterizing the disease and evaluating countermeasures will likely necessitate compliance with the US Food and Drug Administration (FDA) Animal Rule (21 CFR 314.60 for drugs or 21 CFR 601.90 for biological products), as is done for other high consequence pathogens that have high mortality rates and sporadic outbreaks [14–16]. Approval via this mechanism relies on animal model(s) that are well characterized and adequate for showing efficacy, and that adequately recapitulate human disease and efficacy endpoints [14–16]. Such models exist for EBOV and were important for approval of existing vaccines and therapies [17, 18]. SUDV has been characterized in ferrets [19, 20], guinea pigs [21], and via aerosol route in NHPs [22]. However, fewer models exist for SUDV than EBOV and a standardized, well characterized NHP model is still needed [23].

The primary objective of this study was to characterize the disease course in cynomolgus macaques exposed intramuscularly (IM) to SUDV (Gulu variant) to determine if infection in this species is a reproducible and relevant model for the evaluation of filovirus countermeasures under the FDA Animal Rule [24]. Eighteen macaques were exposed IM to a target dose of 1,000 plaque forming units (PFU) of SUDV, at animal biosafety level 4 (ABSL-4). Animals were randomized by sex and body weight to a scheduled euthanasia study arm (n=10) to characterize progression of viremia and clinical pathology, immunology, gross pathology, and histopathology changes at Days 2, 3, 5, 7, and 9 post-exposure (PE), and to a survival study arm (n=8) to confirm the exposure dose was lethal and to measure physiology, clinical pathology, and telemetry kinetics. A third group (n=2) was mock-exposed to sterile phosphate buffered saline (PBS) as controls. There were statistically significant differences between SUDV-exposed animals and mock-exposed animals for time from exposure to onset of abnormality of 26 parameters, including telemetry body temperature, clinical chemistry parameters, hematology parameters, coagulation parameters (activated partial thromboplastin time), viremia (in serum), and serologic biomarkers.

## Results

### Mortality

Eighteen animals were exposed via intramuscular (IM) inoculation with 0.5 mL from a dilution of SUDV stock material corresponding to 1,000 PFU. The delivered dose confirmed by back titration was 1,630 PFU. The two mock-exposed animals inoculated via the IM route with 0.5 mL sterile saline remained healthy throughout the study and were euthanized as planned at the scheduled end of project (EOP) on Day 21 post-exposure. All 10 animals in the serial euthanasia (SE) group survived to their assigned euthanasia days (n=2 each on Day 2, 3, 5, 7, and 9). Seven of eight animals in the survival and physiology time kinetics (SP-TK) group met endpoint criteria and were euthanized: one on each of Days 7, 11, and 13 post-exposure, and two on each of Days 8 and 9 post-exposure. One animal survived to scheduled EOP, Day 21 post-exposure. Kaplan-Meier median time to death for the SUDV-exposed animals in the SP-TK group was 8.99 days (Figure 1A). Mortality is summarized in Table 1.

**Figure 1.**
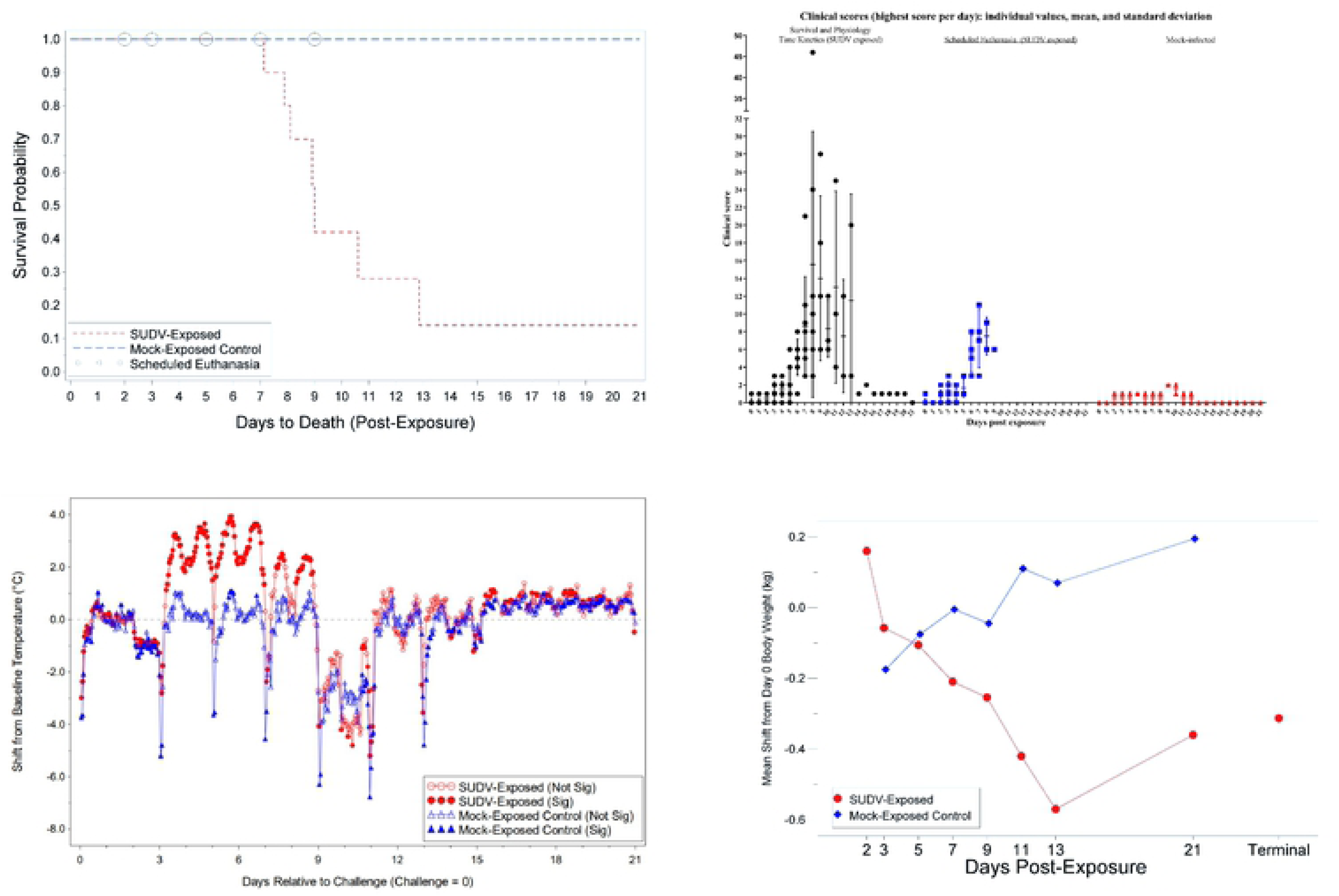
Disease Progression in Cynomolgus Macaques Exposed to SUDV. Group comparisons are displayed for A) Survival probability. B) Daily clinical scores (highest score per day). C) Body temperature changes (°C), mean shift from Day 0, telemetry. D) Body weight change (kg), mean shift from Day 0; T represents Terminal (data collected from animals that met euthanasia criteria were combined and reported as a single terminal time point).

**Table 1.** Mortality Summary.

| Animal ID | Sex | Group Description | Day of death | Final Clinical Score |
| --- | --- | --- | --- | --- |
| 478 | F | SP-TK, Mock-Exposed | Day 21 (EOP) | 0 |
| 495 | M | SP-TK, Mock-Exposed | Day 21 (EOP) | 0 |
| 483 | F | SE | Day 2 | 0 |
| 491 | M | SE | Day 2 | 0 |
| 481 | F | SE | Day 3 | 1 |
| 490 | M | SE | Day 3 | 0 |
| 484 | F | SE | Day 5 | 3 |
| 493 | M | SE | Day 5 | 1 |
| 477 | F | SE | Day 7 | 11 |
| 485 | M | SE | Day 7 | 3 |
| 476 | F | SE | Day 9 | 5 |
| 486 | M | SE | Day 9 | 6 |
| 479 | F | SP-TK | Day 7 | 21 |
| 482 | F | SP-TK | Day 8 | 46 |
| 492 | M | SP-TK | Day 8 | 24 |
| 474 | F | SP-TK | Day 9 | 18 |
| 488 | M | SP-TK | Day 9 | 28 |
| 480 | F | SP-TK | Day 11 | 25 |
| 494 | M | SP-TK | Day 13 | 20 |
| 487 | M | SP-TK | Day 21 (EOP) | 0 |

F – Female; M – Male; EOP – End of Project; SE - Scheduled Euthanasia; SP-TK - Survival and Physiology Time Kinetics

### Clinical Progression of SUDV Disease

#### Clinical Scores

Clinical scores were generated for each animal using thirteen different parameters with assigned numerical scores [18]. The clinical parameters included feed, enrichment, and fluid consumption; stool output; hydration status; hair coat and stool appearance; rectal body temperature changes; body weight changes; presence of nasal discharge, bleeding, or petechia; respiration; and responsiveness. Median and individual highest daily clinical scores for mock-exposed and SUDV-exposed groups are presented in Figure 1B.

Before virus exposure, clinical scores were 3 or less for all 20 animals, with occasional scores for reduced feed consumption or stool output. After the exposure day, the two mock-exposed animals remained healthy and did not exhibit scores greater than 2, associated with slightly diminished activity and sporadic stool abnormalities. At the scheduled euthanasia, the mock-exposed animal clinical scores were 0.

In virus-exposed animals, clinical signs warranting a clinical score were observed infrequently for the first four days following SUDV exposure, with a few sporadic scores for temperature changes, reduced feed consumption or fluid intake, and stool abnormalities. Signs indicative of SUDV disease including diminished activity or responsiveness and mild petechia were first observed on Day 5, with a corresponding increase in clinical scores. The severity of symptoms progressed beginning on Day 6, with most remaining animals exhibiting diminished activity or reduced responses to external stimuli, reduced intake in feed, enrichment and/or fluid, and reduced stool output. By Day 7, all remaining animals exhibited slightly diminished activity or reduced responses to external stimuli, 75% (9 of 12) of animals exhibited mild or moderate petechia, and three animals exhibited bleeding at a site other than the blood collection site.

Between Days 8 and 13, remaining SUDV-exposed animals continued to exhibit diminished activity, reduced responsiveness, petechia and bleeding disorders, reduced feed and enrichment consumption, significant changes in rectal temperature, and stool abnormalities.

The surviving SUDV-exposed animal (Animal 487) had peak clinical scores of 5-6 on Days 6 through 10 but did not exhibit petechia, bleeding, or decreased responsiveness observed in moribund animals, and exhibited clinical scores of two or less for the remainder of the study period. Impaired use of the right arm was observed on Day 7 PE. Clinical examination on of this animal on Day 7 PE revealed several findings. A 6 cm soft subcutaneous mass was observed on the upper abdomen, which resolved by the scheduled necropsy. An area of cutaneous red discoloration with thickening of the skin and firm subjacent skeletal muscle on the right arm at the exposure site was observed during sedation on Day 9 PE; the right axillary lymph node was enlarged. Abnormal observations at the exposure site persisted to the terminal necropsy, consisting of thickened, firm skin and subjacent tissues, resolution of red discoloration, and presence of an overlying cutaneous scab. The nature of the large subcutaneous mass on the abdomen was undetermined, but was considered to likely represent a hematoma due to the soft, fluctuant texture and resolution over the course of the study.

#### Body Temperature

Changes in individual body temperature collected from sedated animals by rectal thermometer were determined by comparing the baseline (average temperature from Day -24 [day of animal transfer into ABSL-4] and Day 0 [prior to SUDV exposure]) with temperatures collected at each scheduled timepoint. For mock-exposed animals, the change from baseline ranged from -2.5°F for animal 478 on Day 7 to +1.3°F for animal 495 on Day 21. In comparison, eight SUDV-exposed animals had increased rectal temperatures that warranted a clinical score (>2°F change from baseline) on Day 5 and more than one animal scored for increased temperature on each data collection day for the remainder of the study period where these animals were alive. Animals that were euthanized due to moribundity typically had decreased rectal temperatures, with 4 of 7 animals exhibiting temperature decreases of >10% from baseline. Animal 487 exhibited increased rectal temperature on Days 5 through 11, with a peak fever of 103.8°F on Day 7. The group mean rectal body temperature in the SUDV-exposed group was significantly increased on Days 2 and 5 and significantly decreased at terminal collection (*p* < 0.05), while there were no significant changes in the mock-exposed group.

The group-based shifts from baseline body temperature collected via telemetry are plotted in Figure 1C. The periodic dips in temperature correspond with sedation on data collection days (Days 3, 5, 7, 9, 11, and 13). With fever defined as a mean temperature value greater than 39°C, the mock-exposed animals and SUDV-exposed animals euthanized on Days 2 and 3 did not reach the fever threshold at any time during the study. Conversely, 12 of 14 remaining animals first met the threshold on Day 3, one animal on Day 4, and the final animal on Day 5. Peak temperatures in these animals ranged from 40.22°C to 41.17°C, occurring on Days 4 to 8. In general, animals that met moribund euthanasia criteria had peak temperature 2 to 6 days prior to succumbing. Temperatures significantly increased from baseline were observed in the SUDV-exposed group from approximately Day 3 to Day 9.

#### Body Weight

The mean change in body weight from baseline (Day 0) for each group is plotted in Figure 1D. The mean body weight at Day 0 was not significantly different between the groups (*p* = 0.9568), implying differences in group mean weights at post-exposure timepoints are associated with effects of exposure and not with inherent differences between the groups at baseline. The mock-exposed animals showed a decrease from baseline weight on Day 3 but weights for these animals increased throughout the study period starting on Day 5. In the SUDV- exposed animals, the mean decrease from baseline body weight was statistically significant at data collection timepoints between Days 3 and 13 and at terminal collection (*p* < 0.05), and body weight loss greater than 5% was observed by Day 5 in individual animals. There were significant group effects (*p* < 0.05) on Days 7, 9, and 11.

#### Clinical Pathology

Changes in hematologic, clinical chemistry, and coagulation parameters were monitored during the study and prior to euthanasia. Parameters that are indicative of bleeding and coagulation disorders, including red blood cell (RBC) counts, hemoglobin (HGB), hematocrit (HCT), activated partial thromboplastin time (aPTT), prothrombin time (PT), platelet (PLT) counts, and PLT distribution width (PDW), were significantly impacted by SUDV exposure but were typically not significantly impacted in mock-exposed animals (Figure 3). Significant decreases as a proportion of baseline (*p* < 0.05) were observed for RBC counts, HGB, and HCT on Days 7 and 9 post-SUDV exposure (Figure 2) while only HGB was significantly decreased as a proportion of baseline for mock-exposed animals, on Days 3 and 9. The group effect for all three parameters when comparing results between the mock-exposed and SUDV-exposed groups was significant on Day 9 (*p* = 0.0185, 0.0237, and 0.0113 for RBC, HGB, and HCT, respectively). In addition, reticulocytes (immature RBCs) were significantly increased as a proportion of baseline (*p* < 0.05) for the SUDV-exposed group on Days 3 and 5 and were significantly decreased on Day 7, corresponding to the decrease in RBCs.

**Figure 2.**
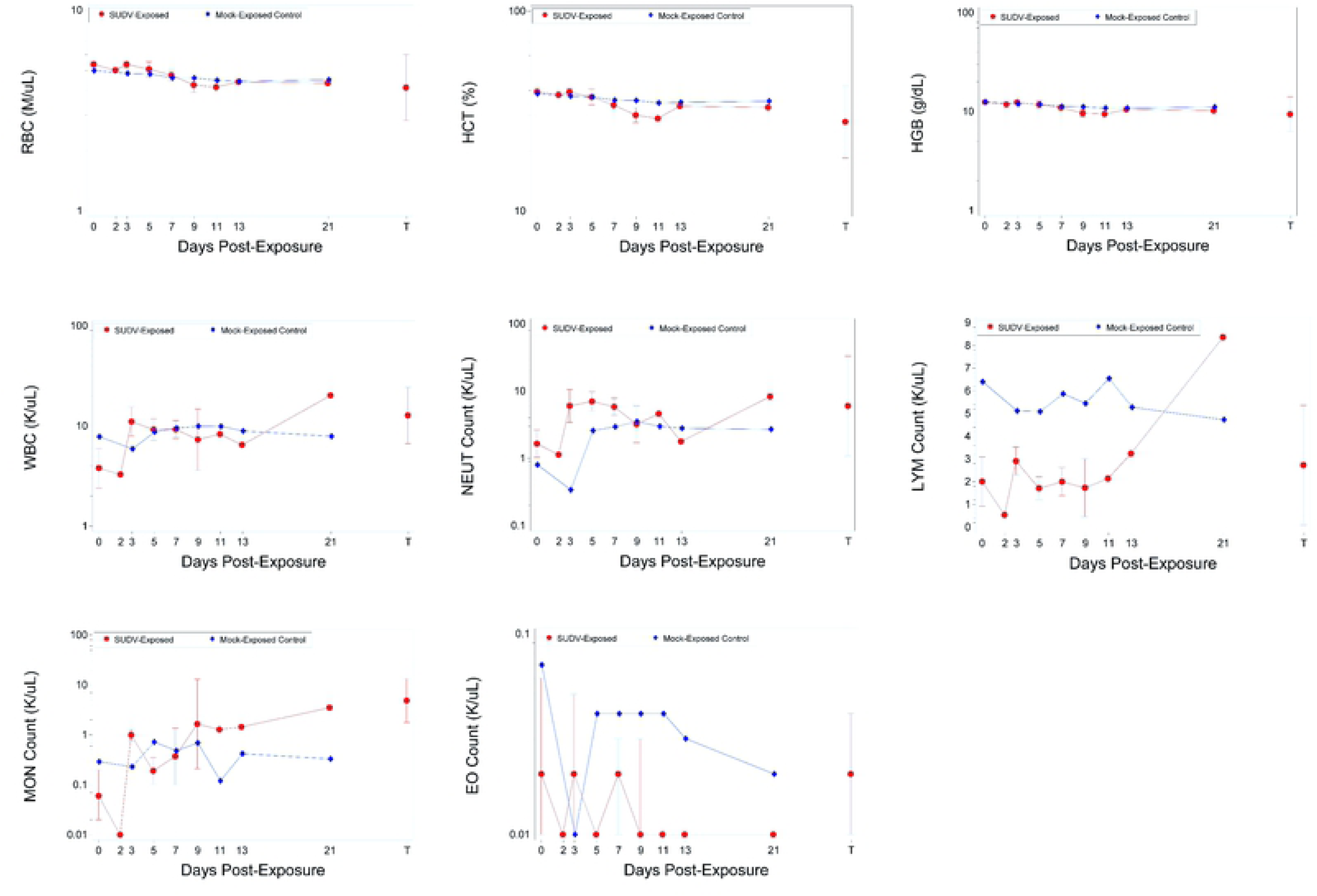
Hematology parameters in SUDV-exposed and mock-exposed cynomolgus macaques. T represents Terminal (data collected from animals that met euthanasia criteria were combined and reported as a single terminal time point). (A) Red blood cell counts. (B) Hematocrit. (C) Hemoglobin. (D) White blood cells. (E) Neutrophils. (F) Lymphocytes. (G) Monocytes. (H) Eosinophils.

**Figure 3.**
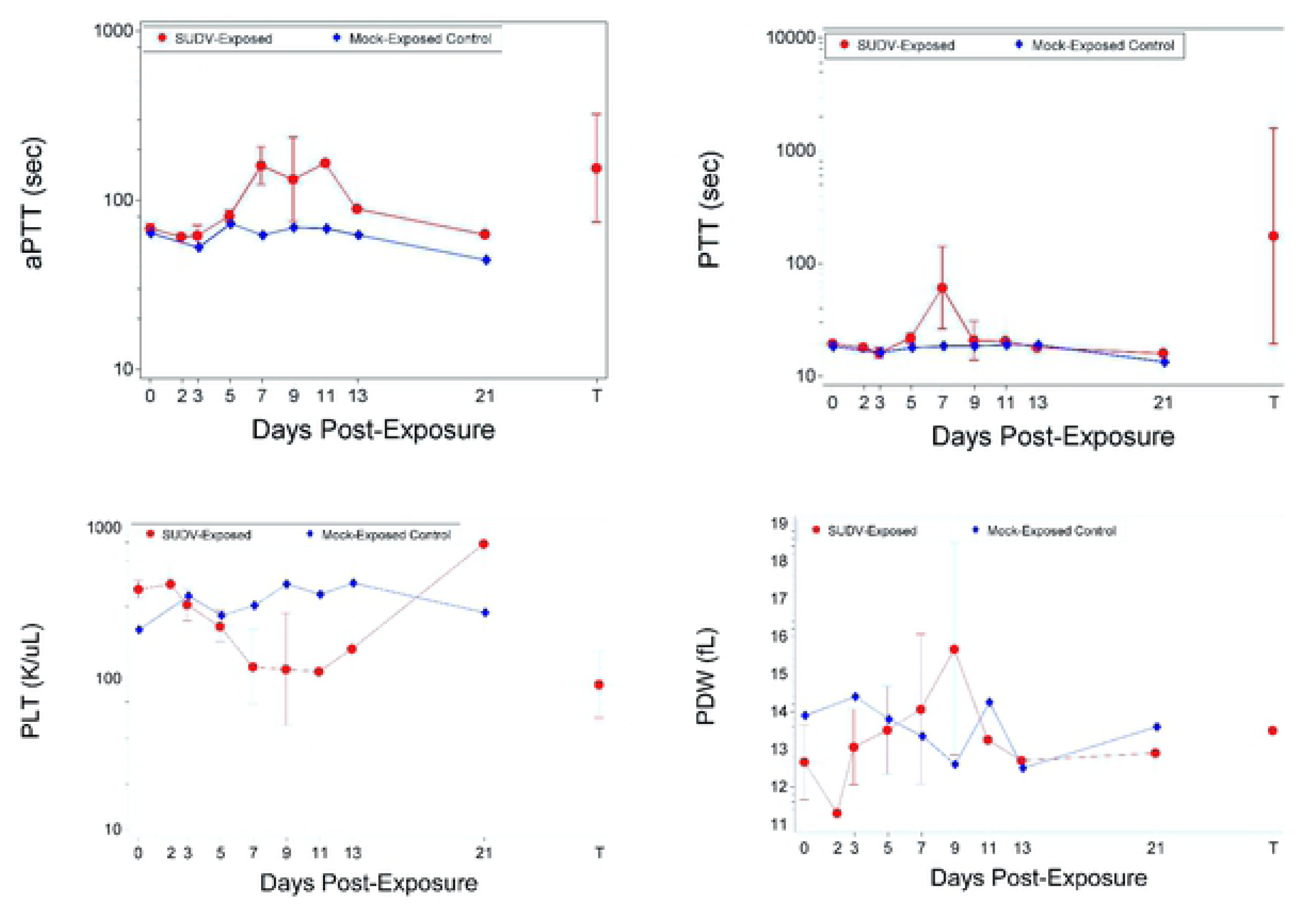
Coagulation parameters in SUDV-exposed and mock-exposed cynomolgus macaques. T represents Terminal (data collected from animals that met euthanasia criteria were combined and reported as a single terminal time point). (A) Activated partial thromboplastin time (aPTT). (B) Prothrombin time (PTT). (C) Platelet counts. (D) Platelet distribution width.

Immune cell populations were also impacted by SUDV exposure but were typically not significantly impacted in mock-exposed animals (Figure 2). For SUDV-exposed animals, there were significant increases as a proportion of baseline (*p* < 0.05) on Days 3, 5, and 7 for white blood cell absolute counts (WBC), neutrophil absolute counts (also increased at terminal; significant group effect on Day 3), and neutrophil relative percentages (mock-exposed animals also exhibited a significant increase on Day 5). Lymphocyte relative percentages were concomitantly decreased on Day 7 and at terminal assessments. However, lymphocyte absolute counts exhibited no significant changes from baseline or group effects. Increased monocyte parameters (absolute counts and relative percentages) also correlated with SUDV disease, but not necessarily with mortality, as the surviving SUDV-exposed animal exhibited these abnormalities. Monocyte absolute counts and relative percentages were significantly increased as a proportion of baseline for SUDV-exposed animals on Day 3 and terminal. Eosinophil absolute counts were significantly decreased (*p* < 0.05) on Days 5, 7, and 13, and eosinophil relative percentages were significantly decreased (*p* < 0.05) on Days 5, 7, and 21; though decreased eosinophils are unlikely to be clinically relevant.

Mock-exposed animals did not exhibit significant changes in coagulation parameters including aPTT, PT, PLT counts, and PDW. SUDV exposure led to significant increases in aPTT as a proportion of baseline (*p* < 0.05) on Days 5, 7, 9, and at terminal collection, significant decreases in PT as a proportion of baseline (*p* < 0.05) on Days 2 and 3 (*p* < 0.05) and significant increases in PT as a proportion of baseline (*p* < 0.05) on Days 5, 7, and terminal, significant decreases in PLT as a proportion of baseline (*p* < 0.05) on Days 3, 5, 7, 9, and terminal, and significant increases in PDW (*p* < 0.05) on Days 5, 7, and 9. The group effect when comparing results between the mock-exposed and SUDV-exposed groups was significant on Study Day 7 for aPTT (*p* = 0.0038), on Days 3, 5, 7, and 9 for PLT (*p* = 0.0321, 0.0013, 0.0306, and 0.0198, respectively), and on Day 9 for PDW (*p* = 0.0115).

Several animals euthanized on or after Day 7 showed increases in clinical chemistry parameters indicative of marked liver damage (Figure 4). Mock-exposed animals only exhibited significant changes for increased total bilirubin (TBIL) on Day 5 and decreased blood urea nitrogen (BUN) on Day 21. For SUDV-exposed animals, there were significant increases (*p* < 0.05) as a proportion of baseline for the following parameters: alanine aminotransferase (ALT) on Days 7, 9, and 11 (significant group effect on Days 7, 9, and 11); alkaline phosphatase (ALP) on Days 2, 3, 7, and 9 (significant group effect on Day 7); bile acids (BA) on Days 5 and 7; gamma glutamyl transferase (GGT) on Day 7 and terminal; total bilirubin (TBIL) on Day 7 (significant group effect on Day 5); and cholesterol (CHOL) on Day 7. There were significant ALT and BA decreases on Day 3.

**Figure 4.**
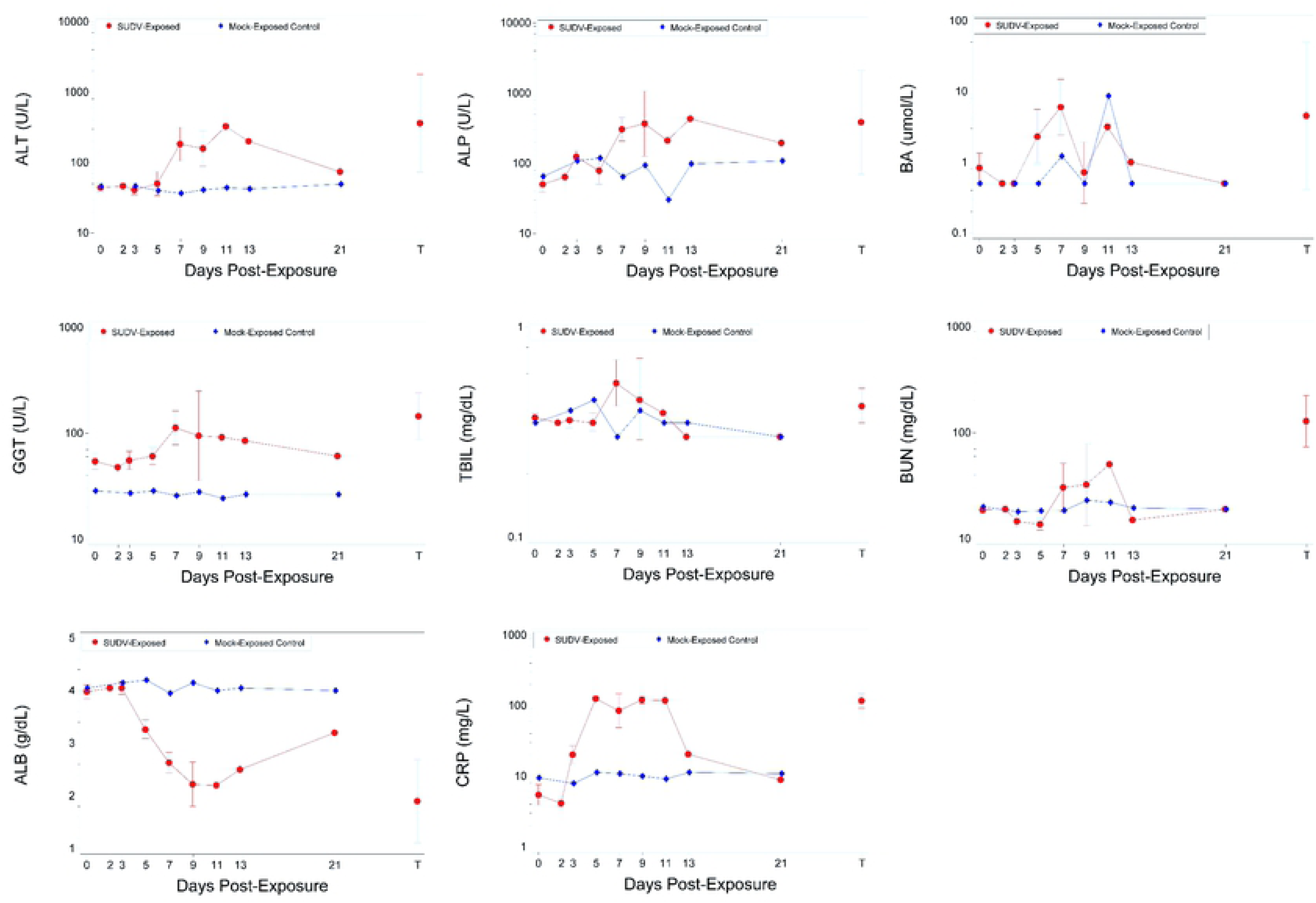
Clinical chemistry parameters in SUDV-exposed and mock-exposed cynomolgus macaques. T represents Terminal (data collected from animals that met euthanasia criteria were combined and reported as a single terminal time point). (A) Alanine aminotransferase. (B) Alkaline phosphatase. (C) Bile acids. (D) Gamma glutamyl transferase. (E) Total bilirubin. (F) Blood urea nitrogen. (G) Albumin. (H) Serum C-reactive protein.

Increased BUN and decreased albumin (ALB) have also been observed in SUDV- exposed animals immediately prior to euthanasia [25]. For BUN, there were significant decreases as a proportion of baseline on Days 3, 5, and 21, and significant increases on Day 7 and terminal; there were no significant group effects. Albumin (ALB) was significantly decreased on Days 5, 7, 9, 11, 13 and terminal (the group effects were significant on Days 5, 7, 9, and 11).

C-reactive protein (CRP), a serum marker for inflammation, was markedly increased in the SUDV-exposed group but remained stable and low in mock-exposed animals (Figure 4). There were significant increases (*p* < 0.05) as a proportion of baseline for the SUDV-exposed group on Days 3, 5, 7, 9, and 11. The group effects were significant on Study Days 5, 7, 9, and 11 (*p* <0.0001, 0.0022, 0.0017, and 0.0317, respectively).

### Virological Progression of SUDV

#### Viremia and Tissue Viral Burden

Viremia was assessed in serum samples collected at scheduled timepoints and at terminal collection and viral burden was assessed in various tissues collected at necropsy. Levels of infectious SUDV in serum and tissues were measured by plaque assay and levels of SUDV genomic RNA equivalents (GE) in serum and tissues were measured by qRT-PCR targeting a region of the glycoprotein gene.

Mock-exposed animals did not have detectable viremia as measured by plaque assay or qRT-PCR at any timepoint tested. In SUDV-exposed animals, the kinetics of infectious virus and viral RNA in serum were similar, with both detected first on Day 3, peak titers were on Day 5 to Day 7, and all remaining SUDV-exposed animals had detectable levels of infectious virus from Day 5 to Day 11 (Figure 5). For most animals, the highest infectious titers were 10^6^ to 10^8^ PFU/mL (peak 1.08 x 10^8^ PFU/mL for animal 493 on Day 5), and peak RNA levels were 10^7^ to 10^10^ GE/mL (peak 1.38 x 10^10^ GE/mL for animal 492 on Day 7). Animal 487 that survived SUDV exposure exhibited low levels of infectious virus in serum on Days 5, 7, and 9 (peak on Day 7 with 1.45 x 10^3^ PFU/mL) and low levels of viral RNA in serum on Days 3 through 13 (peak on Day 9 with 2.11 x 10^5^ GE/mL).

**Figure 5.**
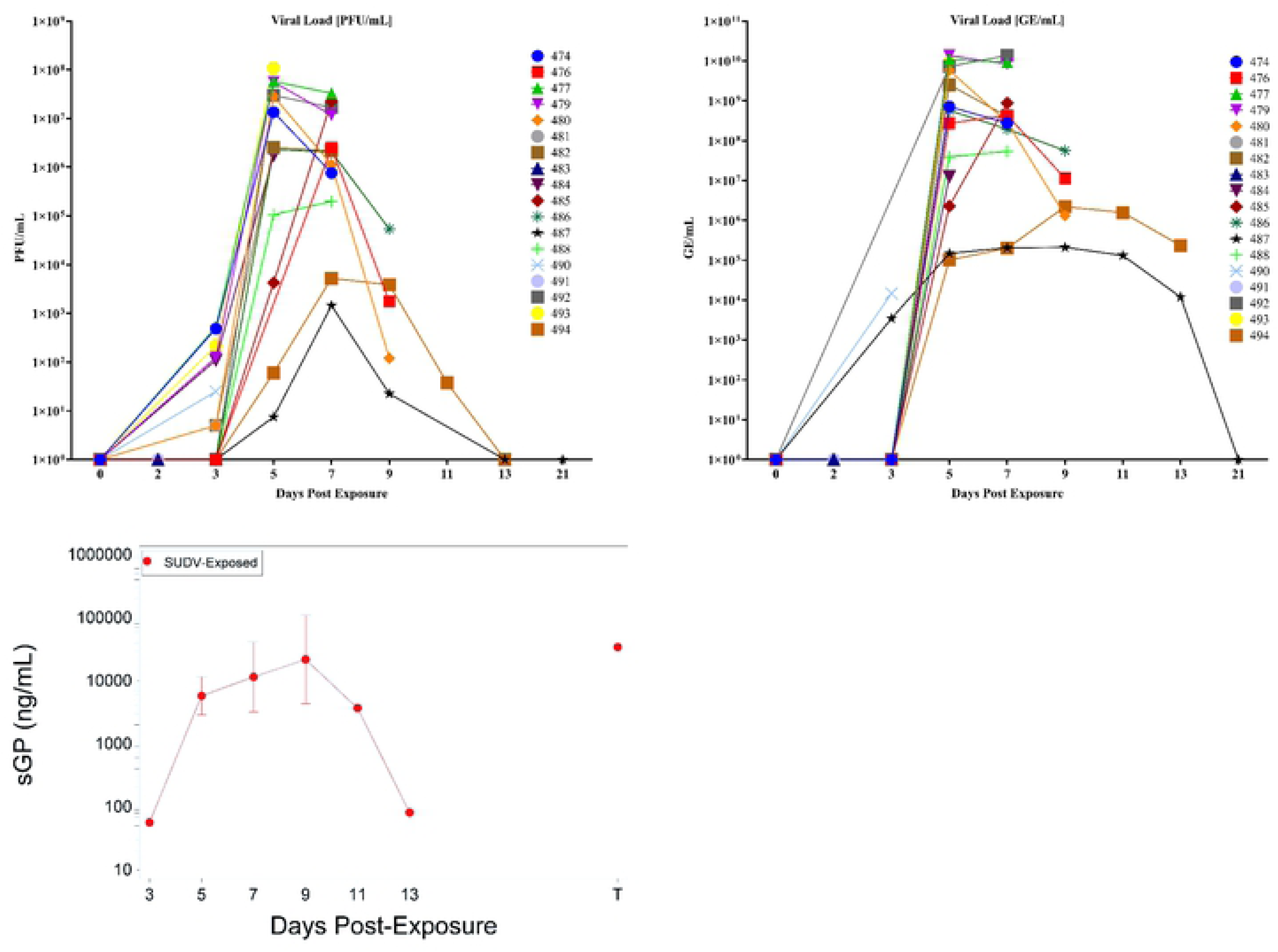
Viral Load. (A) PFU/mL. (B) GE/mL. (C) sGP; T represents Terminal (data collected from animals that met euthanasia criteria were combined and reported as a single terminal time point).

Viral tissue burdens were also assessed by plaque assay and RT-qPCR. Mock-exposed animals did not have detectable infectious SUDV or viral RNA in any of the tissues collected at necropsy on Day 21. The kinetics of viral tissue burden in SUDV-exposed animals were consistent with viremia: infectious virus and viral RNA were detected in some tissues of animals euthanized on schedule on Days 2 and 3, most collected tissues had detectable infectious virus and viral RNA by Day 5, and the highest viral burdens were typically observed in animals euthanized on Days 5 through 9. The largest viral loads (greater than 10^7^ PFU/g and 10^6^-10^8^ GE/µg total RNA) were observed in spleen, liver, gastrointestinal tissues (stomach, duodenum, jejunum, ileum, rectum, and colon), axillary and mediastinal lymph nodes, adrenal gland, and exposure site (skin and underlying subcutis and muscle).

Infectious virus titers in the gastrointestinal (GI) tract tissues tested (stomach, duodenum, jejunum, ileum, rectum, and colon), are shown in Table 2. For animals euthanized on Day 2, animal 483 exhibited low levels of virus in the jejunum and ileum, and animal 491 exhibited no detectable infectious virus in the GI tract. For animals euthanized on Day 3, 481 exhibited low levels of virus in the jejunum and animal 490 exhibited no detectable infectious virus in the GI tract. All animals euthanized on Day 5 and after exhibited detectable virus in all the GI tissues tested (range 8.26 x 10^1^ to 5.41 x 10^7^ PFU/g) except for surviving animal 487 which had no detectable virus in any GI tract tissues on Day 21.

**Table 2.** Viral Load in Gastrointestinal Tract Tissues as Determined by Plaque Assay (PFU/g)

| Animal ID | Group Description | Day of death | Stomach | Duodenum | Jejunum | Ileum | Rectum | Colon |
| --- | --- | --- | --- | --- | --- | --- | --- | --- |
| 478 | SP-TK, Mock-exposed | S, Day 21 | BDL | BDL | BDL | BDL | BDL | BDL |
| 495 | SP-TK, Mock-exposed | S, Day 21 | BDL | BDL | BDL | BDL | BDL | BDL |
| 487 | SP-TK | S, Day 21 | BDL | BDL | BDL | BDL | BDL | BDL |
| 483 | SE | S, Day 2 | BDL | BDL | 4.12 x 10<sup>1</sup> | 1.16 x 10<sup>4</sup> | BDL | BDL |
| 491 | SE | S, Day 2 | BDL | BDL | BDL | BDL | BDL | BDL |
| 481 | SE | S, Day 3 | BDL | BDL | 5.78 x 10<sup>1</sup> | BDL | BDL | BDL |
| 490 | SE | S, Day 3 | BDL | BDL | BDL | BDL | BDL | BDL |
| 484 | SE | S, Day 5 | 8.26 x 10<sup>1</sup> | 4.19 x 10<sup>2</sup> | 1.46 x 10<sup>5</sup> | 1.56 x 10<sup>5</sup> | 3.82 x 10<sup>4</sup> | 7.69 x 10<sup>4</sup> |
| 493 | SE | S, Day 5 | 1.51 x 10<sup>6</sup> | 3.71 x 10<sup>6</sup> | 2.16 x 10<sup>3</sup> | 7.95 x 10<sup>6</sup> | 1.28 x 10<sup>6</sup> | 1.99 x 10<sup>6</sup> |
| 477 | SE | S, Day 7 | 5.06 x 10<sup>6</sup> | 1.09 x 10<sup>7</sup> | 1.51 x 10<sup>7</sup> | 7.63 x 10<sup>6</sup> | 2.63 x 10<sup>7</sup> | 1.52 x 10<sup>7</sup> |
| 485 | SE | S, Day 7 | 1.79 x 10<sup>5</sup> | 8.89 x 10<sup>5</sup> | 4.03 x 10<sup>6</sup> | 5.36 x 10<sup>6</sup> | 4.64 x 10<sup>5</sup> | 1.68 x 10<sup>6</sup> |
| 476 | SE | S, Day 9 | 4.63 x 10<sup>7</sup> | 4.00 x 10<sup>6</sup> | 3.40 x 10<sup>7</sup> | 3.71 x 10<sup>7</sup> | 2.84 x 10<sup>6</sup> | 5.87 x 10<sup>6</sup> |
| 486 | SE | S, Day 9 | 6.22 x 10<sup>6</sup> | 5.29 x 10<sup>7</sup> | 6.93 x 10<sup>6</sup> | 1.05 x 10<sup>7</sup> | 5.40 x 10<sup>6</sup> | 6.38 x 10<sup>6</sup> |
| 479 | SP-TK | US, 7 | 1.48 x 10<sup>6</sup> | 8.58 x 10<sup>6</sup> | 1.80 x 10<sup>7</sup> | 1.08 x 10<sup>7</sup> | 1.13 x 10<sup>7</sup> | 6.86 x 10<sup>6</sup> |
| 482 | SP-TK | US, 8 | 6.39 x 10<sup>6</sup> | 8.85 x 10<sup>6</sup> | 2.65 x 10<sup>7</sup> | 8.26 x 10<sup>6</sup> | 3.38 x 10<sup>7</sup> | 3.26 x 10<sup>7</sup> |
| 492 | SP-TK | US, 8 | 1.70 x 10<sup>5</sup> | 2.56 x 10<sup>6</sup> | 1.78 x 10<sup>7</sup> | 1.69 x 10<sup>6</sup> | 5.95 x 10<sup>6</sup> | 9.26 x 10<sup>6</sup> |
| 474 | SP-TK | US, 9 | 1.61 x 10<sup>7</sup> | 3.03 x 10<sup>6</sup> | 2.42 x 10<sup>7</sup> | 3.45 x 10<sup>7</sup> | 2.73 x 10<sup>7</sup> | 5.16 x 10<sup>7</sup> |
| 488 | SP-TK | US, 9 | 1.01 x 10<sup>6</sup> | 1.13 x 10<sup>5</sup> | 1.06 x 10<sup>6</sup> | 5.31 x 10<sup>6</sup> | 2.88 x 10<sup>6</sup> | 7.16 x 10<sup>6</sup> |
| 480 | SP-TK | US, 11 | 5.41 x 10<sup>7</sup> | 6.14 x 10<sup>6</sup> | 4.14 x 10<sup>7</sup> | 3.59 x 10<sup>7</sup> | 3.58 x 10<sup>7</sup> | 3.48 x 10<sup>7</sup> |
| 494 | SP-TK | US, 13 | 1.47 x 10<sup>5</sup> | 9.64 x 10<sup>5</sup> | 8.86 x 10<sup>6</sup> | 6.82 x 10<sup>6</sup> | 7.68 x 10<sup>6</sup> | 1.07 x 10<sup>7</sup> |

SP-TK—Survival and Pathology Time Kinetics; SE – Serial Sample and Euthanasia; S – Scheduled; US – Unscheduled; BDL - below detection limit (5 PFU/g)

The spleen was also a site for high infectious virus loads (Table 3), with first detection at Day 3 and peak titers of 10^8^ PFU/g observed on Day 5. Infectious virus titers from other tissues (lung, right axillary lymph node, adrenal gland, heart, exposure site, right inguinal lymph node, and mediastinal lymph node) are shown in Table 4. Virus was first detected in these tissues on Day 3, at the exposure site. By Day 5, animal 493 exhibited detectable levels of infectious virus in all these tissues and animal 484 exhibited detectable levels of infectious virus in all tissues except lung. Animals euthanized on Days 7 to 11 exhibited detectable levels of infectious virus in all these tissues (range 4.57 x 10^4^ to 4.90 x 10^7^ PFU/g), with similar titers seen between the different tissues. Animal 494, euthanized on Day 13, exhibited detectable levels of infectious virus in all these tissues, except the heart. Surviving animal 487 did not have detectable infectious SUDV in any of these tissues.

**Table 3.** Viral Load in Spleen and Liver as Determined by Plaque Assay (PFU/g)

| **Animal ID** | **Group Description** | **Day of death** | **Anterior Spleen** | **Posterior Medial Spleen** | **Left Lateral Liver** | **Right Medial Liver** |
| --- | --- | --- | --- | --- | --- | --- |
| 478 | SP-TK, Mock-exposed | S, Day 21 | BDL | BDL | BDL | BDL |
| 495 | SP-TK, Mock-exposed | S, Day 21 | BDL | BDL | BDL | BDL |
| 487 | SP-TK | S, Day 21 | BDL | BDL | BDL | BDL |
| 483 | SE | S, Day 2 | BDL | BDL | BDL | BDL |
| 491 | SE | S, Day 2 | BDL | BDL | BDL | BDL |
| 481 | SE | S, Day 3 | $2.73 \times 10^3$ | $2.16 \times 10^3$ | $3.88 \times 10^3$ | $7.75 \times 10^3$ |
| 490 | SE | S, Day 3 | $8.76 \times 10^2$ | $1.23 \times 10^2$ | $9.43 \times 10^1$ | BDL |
| 484 | SE | S, Day 5 | $2.04 \times 10^8$ | $1.86 \times 10^8$ | $7.44 \times 10^7$ | $4.22 \times 10^7$ |
| 493 | SE | S, Day 5 | $8.30 \times 10^4$ | $2.62 \times 10^8$ | $4.35 \times 10^5$ | $6.34 \times 10^6$ |
| 477 | SE | S, Day 7 | $6.59 \times 10^6$ | $8.18 \times 10^6$ | $4.03 \times 10^6$ | $4.02 \times 10^6$ |
| 485 | SE | S, Day 7 | $3.28 \times 10^7$ | $4.61 \times 10^7$ | $1.74 \times 10^7$ | $7.73 \times 10^6$ |
| 476 | SE | S, Day 9 | $4.33 \times 10^5$ | $2.96 \times 10^5$ | $7.43 \times 10^5$ | $6.77 \times 10^5$ |
| 486 | SE | S, Day 9 | $3.15 \times 10^6$ | $2.98 \times 10^6$ | $2.63 \times 10^5$ | $4.08 \times 10^5$ |
| 479 | SP-TK | US, 7 | $9.33 \times 10^6$ | $2.48 \times 10^7$ | $1.51 \times 10^6$ | $1.48 \times 10^6$ |
| 482 | SP-TK | US, 8 | $4.42 \times 10^6$ | $5.24 \times 10^6$ | $4.39 \times 10^5$ | $6.03 \times 10^5$ |
| 492 | SP-TK | US, 8 | $4.14 \times 10^6$ | $2.64 \times 10^6$ | $3.87 \times 10^5$ | $4.33 \times 10^5$ |
| 474 | SP-TK | US, 9 | $4.49 \times 10^5$ | $4.12 \times 10^5$ | $7.21 \times 10^5$ | $4.22 \times 10^5$ |
| 488 | SP-TK | US, 9 | $6.21 \times 10^4$ | $6.48 \times 10^4$ | $3.23 \times 10^5$ | $2.73 \times 10^5$ |
| 480 | SP-TK | US, 11 | $9.86 \times 10^4$ | $4.70 \times 10^5$ | $7.19 \times 10^5$ | $6.50 \times 10^5$ |
| 494 | SP-TK | US, 13 | $2.04 \times 10^4$ | $1.38 \times 10^4$ | $8.02 \times 10^4$ | $1.01 \times 10^5$ |

SP-TK—Survival and Pathology Time Kinetics; SE – Serial Sample and Euthanasia; S – Scheduled; US – Unscheduled; BDL - below detection limit (5 PFU/g).

**Table 4.** Viral Load in Tissues as Determined by Plaque Assay (PFU/g)

| **Animal ID** | **Group Description** | **Day of death** | **Lung** | **Axillary LN** | **Adrenal Gland** | **Heart** | **Exposure Site** | **Right Inguinal LN** | **Hilar LN** |
| --- | --- | --- | --- | --- | --- | --- | --- | --- | --- |
| 478 | SP-TK, Mock-exposed | S, Day 21 | BDL | BDL | BDL | BDL | BDL | BDL | BDL |
| 495 | SP-TK, Mock-exposed | S, Day 21 | BDL | BDL | BDL | BDL | BDL | BDL | BDL |
| 487 | SP-TK | S, Day 21 | BDL | BDL | BDL | BDL | BDL | BDL | BDL |
| 483 | SE | S, Day 2 | BDL | BDL | BDL | BDL | BDL | BDL | BDL |
| 491 | SE | S, Day 2 | BDL | BDL | BDL | BDL | BDL | BDL | BDL |
| 481 | SE | S, Day 3 | BDL | BDL | BDL | BDL | 7.79 x 10<sup>1</sup> | UD | BDL |
| 490 | SE | S, Day 3 | BDL | BDL | BDL | BDL | 1.28 x 10<sup>1</sup> | BDL | BDL |
| 484 | SE | S, Day 5 | BDL | 3.29 x 10<sup>7</sup> | 2.13 x 10<sup>7</sup> | 7.02 x 10<sup>4</sup> | 1.05 x 10<sup>7</sup> | 2.69 x 10<sup>5</sup> | 1.31 x 10<sup>5</sup> |
| 493 | SE | S, Day 5 | 4.17 x 10<sup>4</sup> | 4.92 x 10<sup>5</sup> | 1.71 x 10<sup>5</sup> | 1.61 x 10<sup>7</sup> | 1.77 x 10<sup>5</sup> | 8.49 x 10<sup>6</sup> | 4.15 x 10<sup>7</sup> |
| 477 | SE | S, Day 7 | 6.00 x 10<sup>5</sup> | 7.70 x 10<sup>6</sup> | 8.81 x 10<sup>6</sup> | 3.33 x 10<sup>5</sup> | 4.86 x 10<sup>6</sup> | 2.81 x 10<sup>7</sup> | 3.29 x 10<sup>6</sup> |
| 485 | SE | S, Day 7 | 4.22 x 10<sup>6</sup> | 2.03 x 10<sup>7</sup> | 4.92 x 10<sup>6</sup> | 4.40 x 10<sup>5</sup> | 6.58 x 10<sup>6</sup> | 8.18 x 10<sup>6</sup> | 3.00 x 10<sup>7</sup> |
| 476 | SE | S, Day 9 | 1.27 x 10<sup>6</sup> | 2.40 x 10<sup>6</sup> | 3.06 x 10<sup>6</sup> | 2.20 x 10<sup>6</sup> | 1.01 x 10<sup>7</sup> | 7.53 x 10<sup>6</sup> | 9.89 x 10<sup>6</sup> |
| 486 | SE | S, Day 9 | 1.51 x 10<sup>6</sup> | 8.40 x 10<sup>5</sup> | 2.40 x 10<sup>6</sup> | 1.10 x 10<sup>6</sup> | 2.39 x 10<sup>7</sup> | 1.79 x 10<sup>6</sup> | 1.95 x 10<sup>6</sup> |
| 479 | SP-TK | US, 7 | 4.50 x 10<sup>6</sup> | 1.41 x 10<sup>5</sup> | 5.52 x 10<sup>6</sup> | 1.71 x 10<sup>6</sup> | 1.01 x 10<sup>7</sup> | 1.40 x 10<sup>6</sup> | 1.17 x 10<sup>7</sup> |
| 482 | SP-TK | US, 8 | 2.38 x 10<sup>5</sup> | 3.98 x 10<sup>6</sup> | 8.24 x 10<sup>5</sup> | 4.57 x 10<sup>4</sup> | 4.90 x 10<sup>7</sup> | 3.94 x 10<sup>6</sup> | 3.76 x 10<sup>5</sup> |
| 492 | SP-TK | US, 8 | 1.06 x 10<sup>6</sup> | 2.37 x 10<sup>6</sup> | 2.19 x 10<sup>7</sup> | 7.31 x 10<sup>5</sup> | 6.79 x 10<sup>5</sup> | 1.47 x 10<sup>5</sup> | 2.37 x 10<sup>6</sup> |
| 474 | SP-TK | US, 9 | 3.52 x 10<sup>6</sup> | 5.32 x 10<sup>6</sup> | 7.02 x 10<sup>6</sup> | 1.47 x 10<sup>6</sup> | 7.34 x 10<sup>6</sup> | 2.21 x 10<sup>6</sup> | 9.16 x 10<sup>6</sup> |
| 488 | SP-TK | US, 9 | 5.33 x 10<sup>5</sup> | 7.40 x 10<sup>4</sup> | 1.64 x 10<sup>5</sup> | 6.10 x 10<sup>4</sup> | 4.97 x 10<sup>6</sup> | 5.81 x 10<sup>6</sup> | 1.38 x 10<sup>5</sup> |
| 480 | SP-TK | US, 11 | 2.33 x 10<sup>6</sup> | 6.60 x 10<sup>6</sup> | 7.82 x 10<sup>6</sup> | 6.96 x 10<sup>5</sup> | 1.92 x 10<sup>7</sup> | 1.33 x 10<sup>7</sup> | 2.67 x 10<sup>6</sup> |
| 494 | SP-TK | US, 13 | 1.29 x 10<sup>6</sup> | 7.20 x 10<sup>5</sup> | 1.11 x 10<sup>5</sup> | BDL | 5.89 x 10<sup>6</sup> | 1.70 x 10<sup>5</sup> | 1.19 x 10<sup>5</sup> |

SP-TK/CP-TK—Survival and Clinical Pathology Time Kinetics; SE – Serial Sample and Euthanasia; LN – lymph node; BDL - below detection limit (5 PFU/g); UD – Undetermined; S, scheduled; US, unscheduled.

Viral GE loads in tissues generally followed similar trends to infectious virus except that not all tissues were positive for viral RNA for all animals (Tables 5 and 6). Mock-exposed animals did not exhibit detectable levels of viral RNA in tissue samples. For animals euthanized on Day 2, animal 491 did not exhibit any detectable viral RNA in the tissues tested and animal 483 was positive only in heart (6.73 x 10^2^ PFU/µg of RNA). All animals euthanized after Day 5 were positive for viral RNA in the following tissues: exposure site, lymph nodes, heart, adrenal gland, spleen, liver, and ileum. A majority of animals also had detectable GE in jejunum, colon, and rectal tissues. More than one animal also had detectable GE in stomach, duodenum, and lung. Animal 487 that survived SUDV exposure exhibited detectable levels of viral RNA in 9 of the 15 tissues tested but did not have detectable GE in lung, adrenal gland, stomach, duodenum, jejunum, or rectum.

**Table 5.** Viral Load in Gastrointestinal Tract Tissues as Determined by qRT-PCR Assay (GE/µg total RNA)

| Animal ID | Group Description | Day of death | Stomach | Duodenum | Jejunum | Ileum | Rectum | Colon |
| --- | --- | --- | --- | --- | --- | --- | --- | --- |
| 478 | SP-TK, Mock-exposed | S, Day 21 | BDL | BDL | BDL | BDL | BDL | BDL |
| 495 | SP-TK, Mock-exposed | S, Day 21 | BDL | BDL | BDL | BDL | BDL | BDL |
| 487 | SP-TK | S, Day 21 | BDL | BDL | BDL | 1.24 x 10<sup>1</sup> | BDL | 1.16 x 10<sup>1</sup> |
| 483 | SE | S, Day 2 | BDL | BDL | BDL | BDL | BDL | BDL |
| 491 | SE | S, Day 2 | BDL | BDL | BDL | BDL | BDL | BDL |
| 481 | SE | S, Day 3 | BDL | BDL | BDL | BDL | BDL | BDL |
| 490 | SE | S, Day 3 | BDL | BDL | BDL | BDL | BDL | BDL |
| 484 | SE | S, Day 5 | BDL | 8.47 x 10<sup>3</sup> | 1.86 x 10<sup>3</sup> | 5.85 x 10<sup>3</sup> | 1.33 x 10<sup>4</sup> | BDL |
| 493 | SE | S, Day 5 | BDL | BDL | 8.47 x 10<sup>3</sup> | 2.55 x 10<sup>4</sup> | 4.76 x 10<sup>1</sup> | 2.42 x 10<sup>3</sup> |
| 477 | SE | S, Day 7 | BDL | BDL | 4.30 x 10<sup>1</sup> | 3.98 x 10<sup>5</sup> | 4.62 x 10<sup>3</sup> | 4.43 x 10<sup>3</sup> |
| 485 | SE | S, Day 7 | BDL | BDL | BDL | 2.08 x 10<sup>3</sup> | 6.48 x 10<sup>0</sup>① | 1.06 x 10<sup>1</sup> |
| 476 | SE | S, Day 9 | 6.23 x 10<sup>4</sup> | BDL | BDL | 1.52 x 10<sup>6</sup> | 1.03 x 10<sup>4</sup> | 7.70 x 10<sup>3</sup> |
| 486 | SE | S, Day 9 | 1.21 x 10<sup>4</sup> | 7.56 x 10<sup>2</sup> | 7.54 x 10<sup>3</sup> | 1.43 x 10<sup>5</sup> | 5.58 x 10<sup>4</sup> | 2.20 x 10<sup>5</sup> |
| 479 | SP-TK | US, 7 | BDL | BDL | BDL | 6.81 x 10<sup>4</sup> | BDL | 3.03 x 10<sup>3</sup> |
| 482 | SP-TK | US, 8 | BDL | 9.01 x 10<sup>5</sup> | BDL | 2.08 x 10<sup>4</sup> | BDL | 4.15 x 10<sup>2</sup> |
| 492 | SP-TK | US, 8 | BDL | 7.36 x 10<sup>3</sup> | 3.30 x 10<sup>5</sup> | 5.42 x 10<sup>5</sup> | 6.51 x 10<sup>5</sup> | 4.89 x 10<sup>5</sup> |
| 474 | SP-TK | US, 9 | BDL | 3.96 x 10<sup>3</sup> | 1.14 x 10<sup>1</sup> | 1.37 x 10<sup>6</sup> | 4.42 x 10<sup>5</sup> | 3.70 x 10<sup>5</sup> |
| 488 | SP-TK | US, 9 | BDL | 3.16 x 10<sup>4</sup> | 2.59 x 10<sup>4</sup> | 2.88 x 10<sup>5</sup> | 1.37 x 10<sup>5</sup> | 6.83 x 10<sup>4</sup> |
| 480 | SP-TK | US, 11 | 1.47 x 10<sup>6</sup> | BDL | 1.38 x 10<sup>6</sup> | 1.63 x 10<sup>6</sup> | 1.91 x 10<sup>6</sup> | 9.54 x 10<sup>5</sup> |
| 494 | SP-TK | US, 13 | BDL | 1.17 x 10<sup>6</sup> | BDL | 1.70 x 10<sup>4</sup> | 2.41 x 10<sup>4</sup> | 2.90 x 10<sup>4</sup> |

SP-TK/CP-TK—Survival and Clinical Pathology Time Kinetics; SE – Serial Sample and Euthanasia; BDL - below detection limit (<10 GE/ μg of RNA); ① below lower limit of quantification (10 GE/ μg of RNA); S, scheduled; US, unscheduled.

**Table 6.**
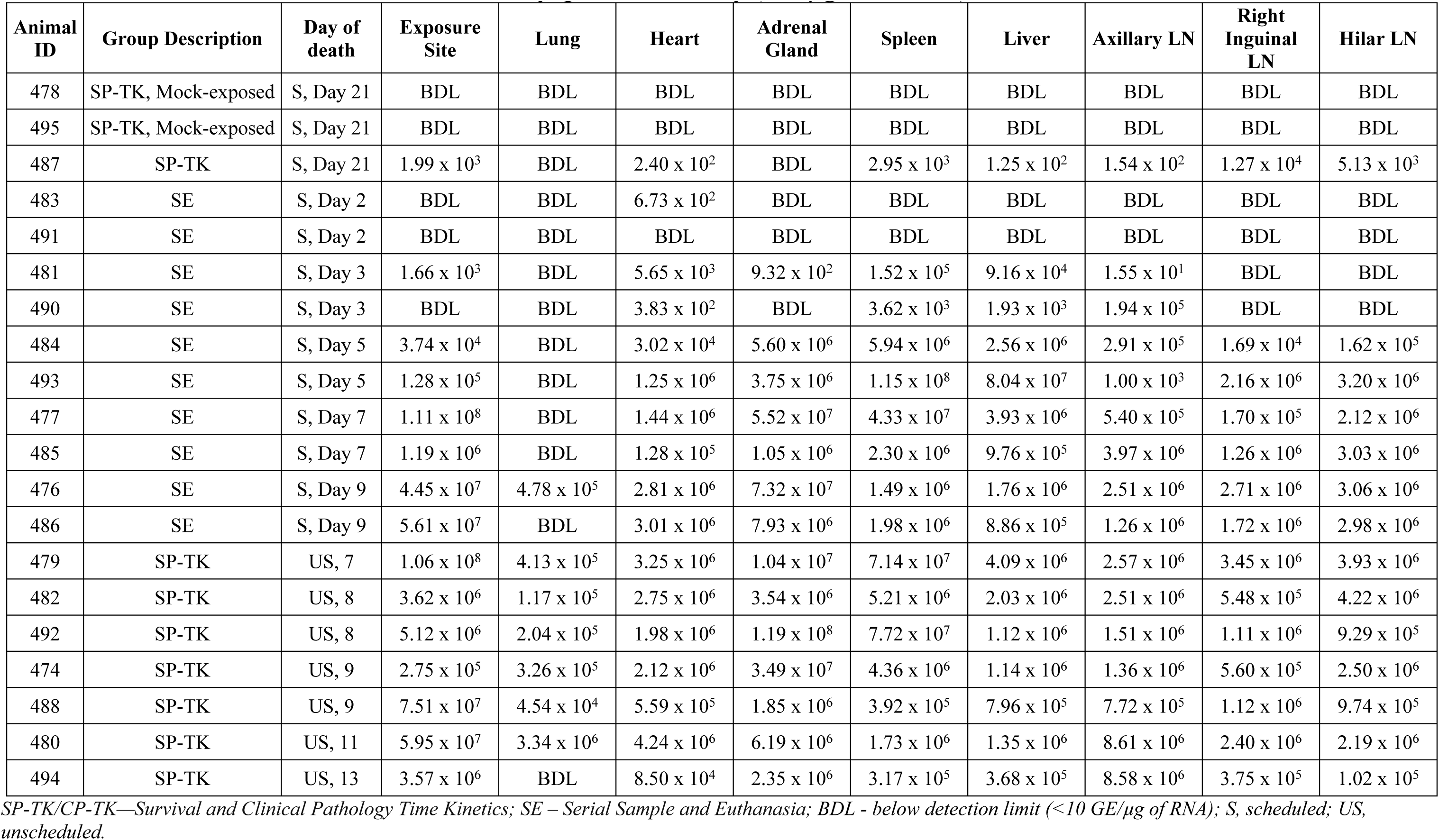
Viral Load in Tissues as Determined by qRT-PCR Assay (GE/µg total RNA)

| Animal ID | Group Description | Day of death | Exposure Site | Lung | Heart | Adrenal Gland | Spleen | Liver | Axillary LN | Right Inguinal LN | Hilar LN |
| --- | --- | --- | --- | --- | --- | --- | --- | --- | --- | --- | --- |
| 478 | SP-TK, Mock-exposed | S, Day 21 | BDL | BDL | BDL | BDL | BDL | BDL | BDL | BDL | BDL |
| 495 | SP-TK, Mock-exposed | S, Day 21 | BDL | BDL | BDL | BDL | BDL | BDL | BDL | BDL | BDL |
| 487 | SP-TK | S, Day 21 | $1.99 \times 10^3$ | BDL | $2.40 \times 10^2$ | BDL | $2.95 \times 10^3$ | $1.25 \times 10^2$ | $1.54 \times 10^2$ | $1.27 \times 10^4$ | $5.13 \times 10^3$ |
| 483 | SE | S, Day 2 | BDL | BDL | $6.73 \times 10^2$ | BDL | BDL | BDL | BDL | BDL | BDL |
| 491 | SE | S, Day 2 | BDL | BDL | BDL | BDL | BDL | BDL | BDL | BDL | BDL |
| 481 | SE | S, Day 3 | $1.66 \times 10^3$ | BDL | $5.65 \times 10^3$ | $9.32 \times 10^2$ | $1.52 \times 10^5$ | $9.16 \times 10^4$ | $1.55 \times 10^1$ | BDL | BDL |
| 490 | SE | S, Day 3 | BDL | BDL | $3.83 \times 10^2$ | BDL | $3.62 \times 10^3$ | $1.93 \times 10^3$ | $1.94 \times 10^5$ | BDL | BDL |
| 484 | SE | S, Day 5 | $3.74 \times 10^4$ | BDL | $3.02 \times 10^4$ | $5.60 \times 10^6$ | $5.94 \times 10^6$ | $2.56 \times 10^6$ | $2.91 \times 10^5$ | $1.69 \times 10^4$ | $1.62 \times 10^5$ |
| 493 | SE | S, Day 5 | $1.28 \times 10^5$ | BDL | $1.25 \times 10^6$ | $3.75 \times 10^6$ | $1.15 \times 10^8$ | $8.04 \times 10^7$ | $1.00 \times 10^3$ | $2.16 \times 10^6$ | $3.20 \times 10^6$ |
| 477 | SE | S, Day 7 | $1.11 \times 10^8$ | BDL | $1.44 \times 10^6$ | $5.52 \times 10^7$ | $4.33 \times 10^7$ | $3.93 \times 10^6$ | $5.40 \times 10^5$ | $1.70 \times 10^5$ | $2.12 \times 10^6$ |
| 485 | SE | S, Day 7 | $1.19 \times 10^6$ | BDL | $1.28 \times 10^5$ | $1.05 \times 10^6$ | $2.30 \times 10^6$ | $9.76 \times 10^5$ | $3.97 \times 10^6$ | $1.26 \times 10^6$ | $3.03 \times 10^6$ |
| 476 | SE | S, Day 9 | $4.45 \times 10^7$ | $4.78 \times 10^5$ | $2.81 \times 10^6$ | $7.32 \times 10^7$ | $1.49 \times 10^6$ | $1.76 \times 10^6$ | $2.51 \times 10^6$ | $2.71 \times 10^6$ | $3.06 \times 10^6$ |
| 486 | SE | S, Day 9 | $5.61 \times 10^7$ | BDL | $3.01 \times 10^6$ | $7.93 \times 10^6$ | $1.98 \times 10^6$ | $8.86 \times 10^5$ | $1.26 \times 10^6$ | $1.72 \times 10^6$ | $2.98 \times 10^6$ |
| 479 | SP-TK | US, 7 | $1.06 \times 10^8$ | $4.13 \times 10^5$ | $3.25 \times 10^6$ | $1.04 \times 10^7$ | $7.14 \times 10^7$ | $4.09 \times 10^6$ | $2.57 \times 10^6$ | $3.45 \times 10^6$ | $3.93 \times 10^6$ |
| 482 | SP-TK | US, 8 | $3.62 \times 10^6$ | $1.17 \times 10^5$ | $2.75 \times 10^6$ | $3.54 \times 10^6$ | $5.21 \times 10^6$ | $2.03 \times 10^6$ | $2.51 \times 10^6$ | $5.48 \times 10^5$ | $4.22 \times 10^6$ |
| 492 | SP-TK | US, 8 | $5.12 \times 10^6$ | $2.04 \times 10^5$ | $1.98 \times 10^6$ | $1.19 \times 10^8$ | $7.72 \times 10^7$ | $1.12 \times 10^6$ | $1.51 \times 10^6$ | $1.11 \times 10^6$ | $9.29 \times 10^5$ |
| 474 | SP-TK | US, 9 | $2.75 \times 10^5$ | $3.26 \times 10^5$ | $2.12 \times 10^6$ | $3.49 \times 10^7$ | $4.36 \times 10^6$ | $1.14 \times 10^6$ | $1.36 \times 10^6$ | $5.60 \times 10^5$ | $2.50 \times 10^6$ |
| 488 | SP-TK | US, 9 | $7.51 \times 10^7$ | $4.54 \times 10^4$ | $5.59 \times 10^5$ | $1.85 \times 10^6$ | $3.92 \times 10^5$ | $7.96 \times 10^5$ | $7.72 \times 10^5$ | $1.12 \times 10^6$ | $9.74 \times 10^5$ |
| 480 | SP-TK | US, 11 | $5.95 \times 10^7$ | $3.34 \times 10^6$ | $4.24 \times 10^6$ | $6.19 \times 10^6$ | $1.73 \times 10^6$ | $1.35 \times 10^6$ | $8.61 \times 10^6$ | $2.40 \times 10^6$ | $2.19 \times 10^6$ |
| 494 | SP-TK | US, 13 | $3.57 \times 10^6$ | BDL | $8.50 \times 10^4$ | $2.35 \times 10^6$ | $3.17 \times 10^5$ | $3.68 \times 10^5$ | $8.58 \times 10^6$ | $3.75 \times 10^5$ | $1.02 \times 10^5$ |

SP-TK/CP-TK—Survival and Clinical Pathology Time Kinetics; SE – Serial Sample and Euthanasia; BDL - below detection limit ( $<10$ GE/ μg of RNA); S, scheduled; US, unscheduled.

#### Soluble Glycoprotein

The quantities of circulating soluble glycoprotein (sGP), a viral protein produced during active SUDV replication, were determined using the SUDV Soluble GP ELISA Kit (IBT Bioservices), which allows for quantitative measurement of SUDV sGP in NHP serum (Figure 5). The following samples were not available for analysis: animal 488, Day 7; animal 492, Days 3 and 7; animal 492, Day 8 (terminal); animal 474 Day 9 (terminal); animal 480, Day 11; animal 482, Day 8; and animal 488, Day 9.

No sGP was detected in samples from mock-exposed animals. For SUDV-exposed animals, kinetics of sGP were similar to detection of infectious virus and viral RNA, with detectable levels of sGP observed in two animals on Day 3, most animals (12 of 14) were positive on Day 5, and all remaining animals for which sample was available (n = 10) were positive on Day 7 with peak sGP concentrations of 10^4^ to 10^5^ ng/mL observed on Days 5 to 7. Survivor animal 487 exhibited measurable sGP levels from Days 7 to 13.

### Immunological Response to SUDV Exposure

#### Cytokine and Chemokine Expression

Collected sera were analyzed for specific cytokines and chemokines using the MILLIPLEX MAP Non-Human Primate Cytokine Magnetic Bead Panel (Figure 6). The following analytes were assessed: Tumor necrosis factor alpha (TNF-α), granulocyte-macrophage colony-stimulating factor (GM-CSF), transforming growth factor alpha (TGF-α), Granulocyte-colony stimulating factor (G-CSF) interferon gamma (IFN-g), interleukin (IL)-2, IL-10, IL-15, IL-17, IL-1ra, IL-13, IL-1b, IL-4, IL-5, IL-6, IL-8, macrophage inflammatory protein (MIP)-1a, monocyte chemotactic protein-1 (MCP-1), MIP-1b, IL-12/23 (p40),vascular endothelial growth factor (VEGF), IL-18, and soluble FD40 ligand (sCD40L; not included in figure as data were not log transformed as with all other parameters).

**Figure 6.**
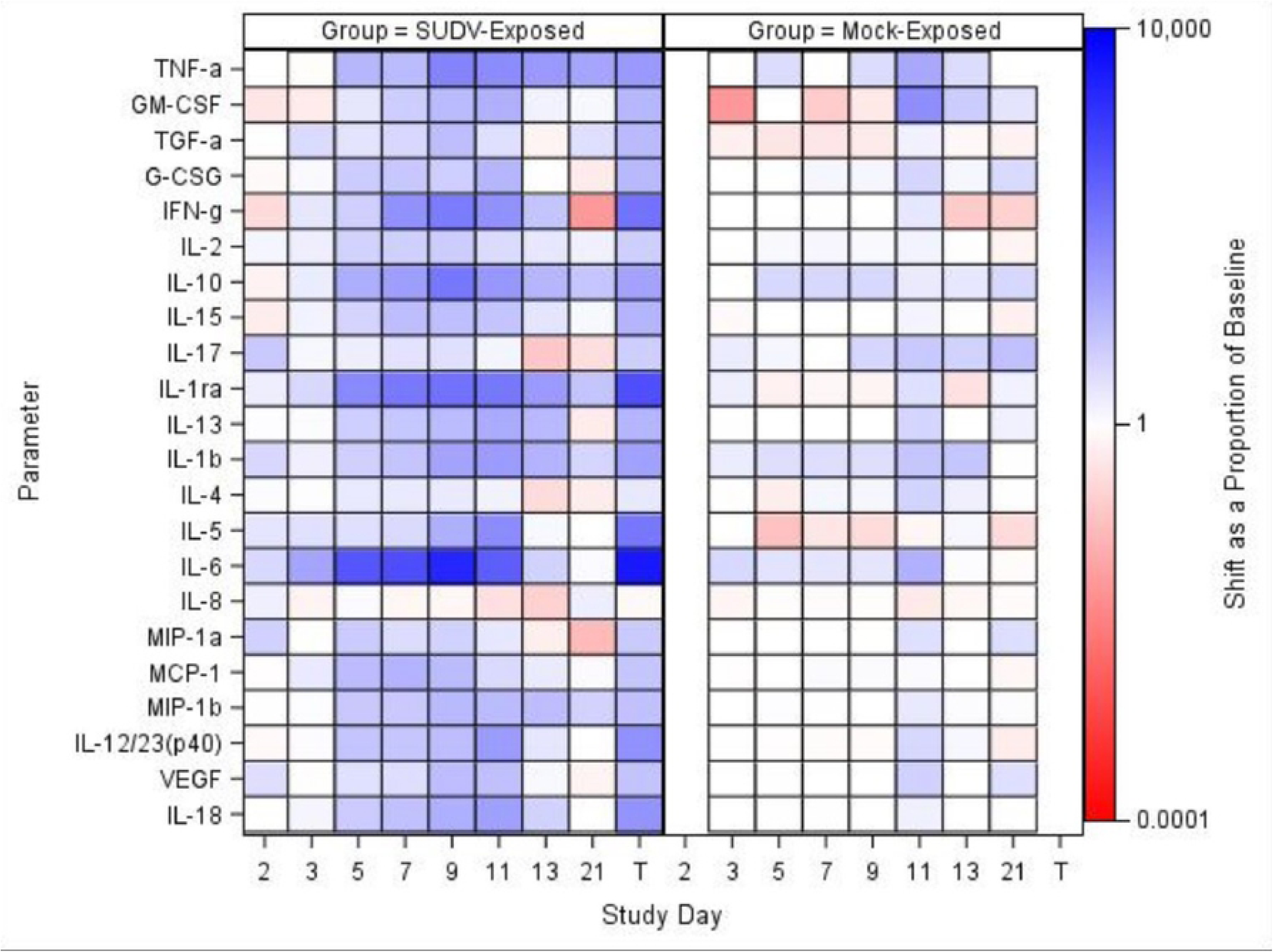
Biomarker heat map; all biomarker parameters except sCD40L were log- transformed; therefore, sCD40L is not included in the heatmap. TNF-α data are not available for mock-exposed animals on Day 21. T represents Terminal (data collected from animals that met euthanasia criteria were combined and reported as a single terminal time point).

For mock-exposed animals, levels of cytokines and chemokines were stable, with sporadic increases in only a few values: On Day 11, IL-4 and TNF-α had significant increases from baseline and on Days 3, 7, and 9, sCD40L had significant increases from baseline. The remaining parameters did not show significant increases or decreases from baseline at any collection timepoint evaluated. Temporally in the SUDV-exposed group, many parameters that may be involved in inflammation, coagulopathy, and endothelial permeability during filovirus infection [26–28], exhibited significant (*p*<0.05) increases from baseline. Some of these parameters were increased as early as Day 3 (TGF-α, IL-2, IL-1ra, IL-5, IL-6, and MCP-1), with the majority being increased on Days 5 through 9. Specifically, a number of interleukins and chemotactic proteins have been observed to be increased in human filovirus infections. Some studies report that lower levels of IL-6, IL-10 (an anti-inflammatory cytokine), IL-8, and MIP-1β may be early predictors of survival as they have been substantially increased in fatal cases [26–28]. In this study, SUDV-exposed animals exhibited increased IL-10 (significant increases as a proportion of baseline for the SUDV-exposed group on Study Days 5, 7, 9, and 11) and IL-6 (significant increases as a proportion of baseline for the SUDV-exposed group on Study Days 3, 5, 7, 9 and at Terminal). IL-1β, IL-5, and IL-18 have also been found elevated in human cases, but are not as correlative with survival. In the SUDV-exposed animals, there were significant increases as a proportion of baseline for: IL-1β (Days 5, 7, and 9), IL-5 (Days 3, 5, 7, and 9), and IL-18 (Days 5, 7, 9, 11 and at Terminal). IL-15, IL-13, and IL-12/23 (p40) were also significantly increased as a proportion of baseline on Days 5, 7, and 9 (IL-15 was increased on Days 11 and 13 as well, and IL-13 at Terminal).

Furthermore, MCP-1, MIP-1α, and MIP-1β—involved in attracting immune cells to the site of inflammation and may be involved in immunopathology during EVD — were increased as well. For MCP-1, there were significant increases as a proportion of baseline for the SUDV- exposed group on Study Days 3, 5, 7, 9, and 11; the group effects were significant on Study Days 7 and 11. For MIP-1α, there were significant increases as a proportion of baseline for the SUDV- exposed group on Study Days 2, 5, and 7; the group effect was significant on Study Day 7.

Finally, for MIP-1β, there were significant increases as a proportion of baseline for the SUDV- exposed group on Study Days 5, 7, 9, 11, and 13; the group effects were significant on Study Days 7 and 9; increases are also observed in fatal human SUDV infections.

More recently, studies in humans and NHPs have shown that high levels of interleukin 1 receptor antagonist (IL-1Ra, an inhibitor of IL1β activity) are found in fatal cases [29]. In this study, there were significant increases as a proportion of baseline for the SUDV-exposed group throughout (Days 3, 5, 7, 9, 11, 21 and Terminal); the group effects were significant on Study Days 5, 7, and 9.

Additionally, while the dataset of human SUDV infections is smaller, previous work has shown that increased TNF-α, IFN-γ, and IL-2 may be less relevant in SVD than EVD [27]. Increased TNFα and IFN- γ have been observed in fatal human cases of EBOV, but the trend is less clear in human SUDV infections. However, in this study, TNFα – which is implicated in inflammation and coagulopathy – exhibited increases. There were significant increases as a proportion of baseline for the SUDV-exposed group on Study Days 5, 7, 9, 11 and at Terminal and a significant increase as a proportion of baseline for the mock-exposed group on Study Day 11. Similarly, IL-2 exhibited significant increases as a proportion of baseline for the SUDV-exposed group on Study Days 3, 5, 7, 9, and 11. The group effects were significant on Study Days 5 and 11. Furthermore, Interferon γ increased in some animals after Day 5 post exposure; IFN α and β are not measured in this panel, but in EVD, Type 1 Interferon responses have been observed to be impaired, which can lead to increased Type II interferon [26]. There were significant increases in IFN- γ as a proportion of baseline for the SUDV-exposed group on Study Days 5, 7, 9, and 11; the group effects were significant on Study Days 7 and 9.

VEGF, G-CSF, and GM-CSF, which are involved in growth differentiation, were also increased. The role of VEGF in filovirus disease remains unclear, but colony-stimulating factors appear to be increased in fatal human cases. In the SUDV-exposed animals, for VEGF, there were significant increases as a proportion of baseline on Study Days 5, 7, and 9; there was no significant group effect on any study day. For G-CSF, there were significant increases as a proportion of baseline for the SUDV-exposed group on Study Days 5, 7, and 9; there was no significant group effect on any study day. Finally, for GM-CSF, there were significant increases as a proportion of baseline for the SUDV-exposed group on Study Days 5, 7 and 9; the group effect was significant on Study Day 7.

Finally, decreased levels of sCD40L were found, which is expected in non-survivors based on human data. High levels of sCD40L have been detected in survivors leading to the suggestion that sCD40L could be a novel biomarker to predict clinical outcome [28]. As with other parameters discussed above, changes were more frequent and more drastic in animals euthanized later in the study. There was a significant increase from baseline for the SUDV- exposed group on Study Day 3, and the group effects were significant on Study Days 7 and 9.

Changes in IL-17 (a pro-inflammatory cytokine) were minor, except for in the surviving SUDV-exposed animal 487; IL-17 in this animal peaked on Day 7 post exposure. The role of IL-17 and Th17 cells in filovirus infections remains an area of investigation [26], but it is possible the spike in IL-17 had a protective effect in this animal.

### Pathologic Progression of SUDV Disease

#### Gross Observations

No significant macroscopic observations were noted in the two mock-exposed animals at the Day 21 PE terminal necropsy. Gross observations in SUDV-exposed animals are summarized in Table 7 and representative images are in Figure 7.

**Figure 7.**
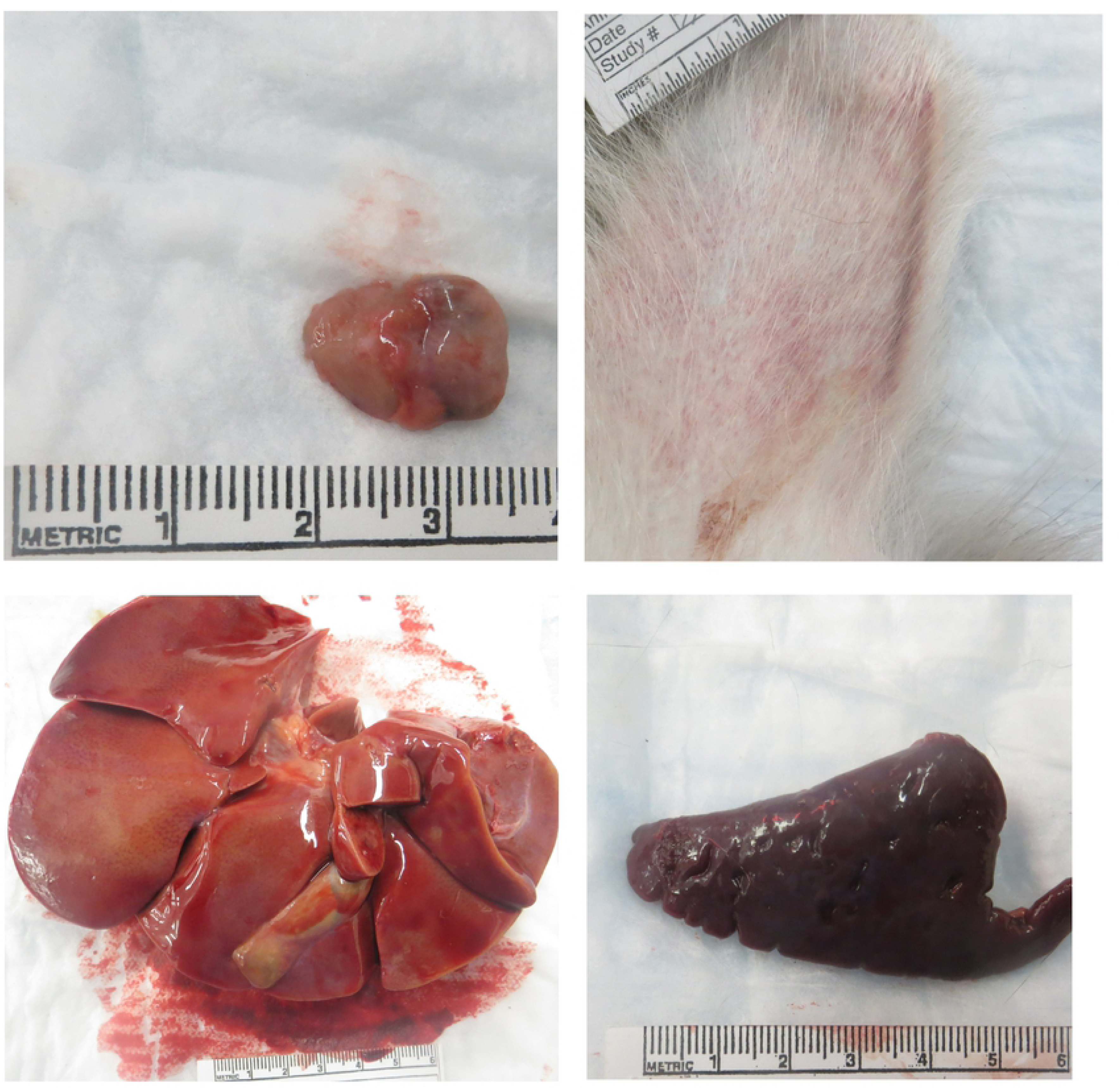
Representative images from gross necropsy. Macroscopic Findings at Necropsy in cynomolgus macaques intramuscularly exposed to SUDV Gulu. (A) Animal 480 (unscheduled euthanasia, Day 11), enlarged axillary lymph node. (B) Animal 494 (unscheduled euthanasia, Day 13), skin rash. (C) Animal 488 (unscheduled euthanasia, Day 9), pale liver. (D) Animal 492 (unscheduled euthanasia, Day 8), enlarged spleen.

**Table 7.**
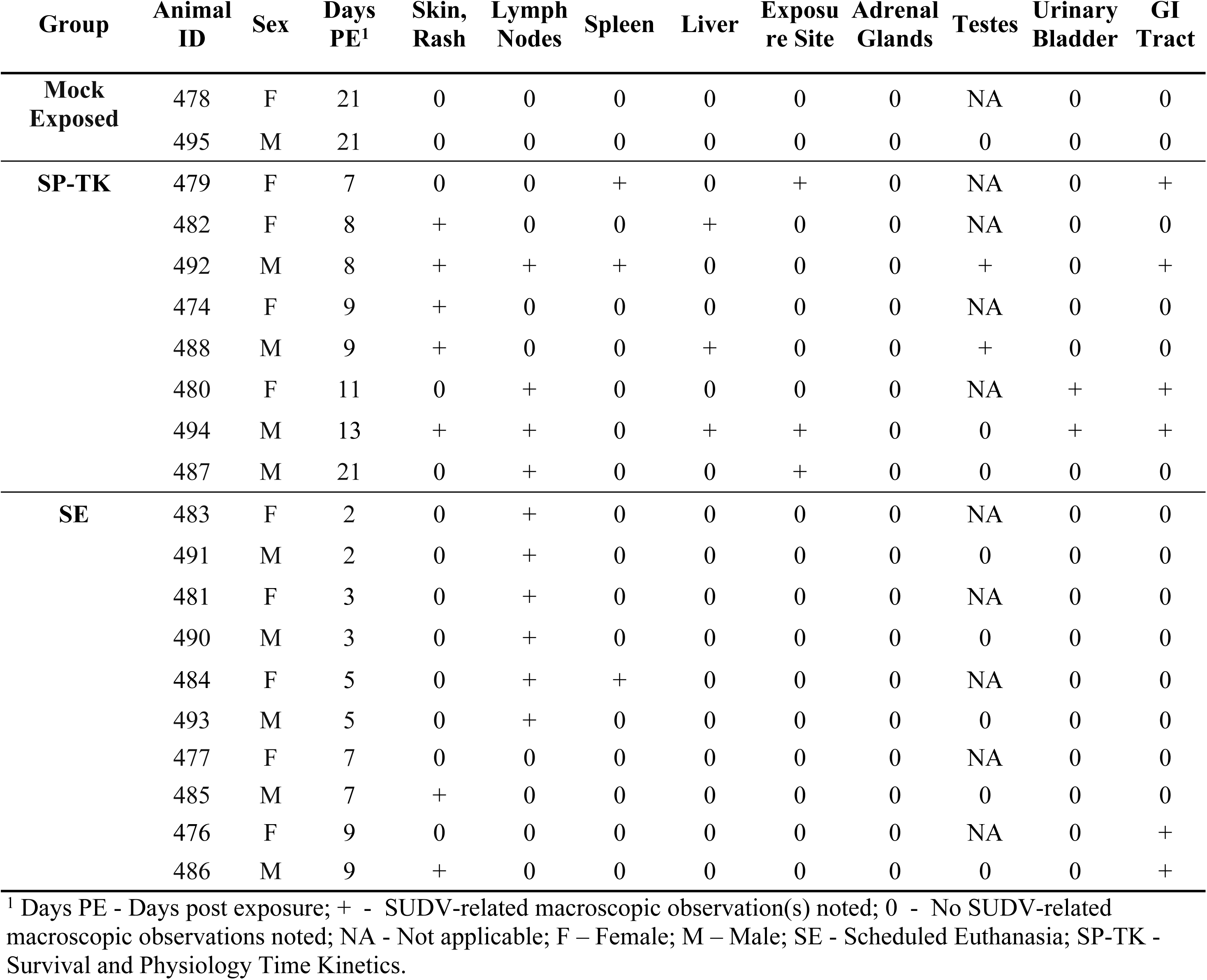
SUDV-related Macroscopic Findings.

| Group | Animal ID | Sex | Days PE<sup>1</sup> | Skin, Rash | Lymph Nodes | Spleen | Liver | Exposure Site | Adrenal Glands | Testes | Urinary Bladder | GI Tract |
| --- | --- | --- | --- | --- | --- | --- | --- | --- | --- | --- | --- | --- |
| **Mock Exposed** | 478 | F | 21 | 0 | 0 | 0 | 0 | 0 | 0 | NA | 0 | 0 |
|  | 495 | M | 21 | 0 | 0 | 0 | 0 | 0 | 0 | 0 | 0 | 0 |
| **SP-TK** | 479 | F | 7 | 0 | 0 | + | 0 | + | 0 | NA | 0 | + |
|  | 482 | F | 8 | + | 0 | 0 | + | 0 | 0 | NA | 0 | 0 |
|  | 492 | M | 8 | + | + | + | 0 | 0 | 0 | + | 0 | + |
|  | 474 | F | 9 | + | 0 | 0 | 0 | 0 | 0 | NA | 0 | 0 |
|  | 488 | M | 9 | + | 0 | 0 | + | 0 | 0 | + | 0 | 0 |
|  | 480 | F | 11 | 0 | + | 0 | 0 | 0 | 0 | NA | + | + |
|  | 494 | M | 13 | + | + | 0 | + | + | 0 | 0 | + | + |
|  | 487 | M | 21 | 0 | + | 0 | 0 | + | 0 | 0 | 0 | 0 |
| **SE** | 483 | F | 2 | 0 | + | 0 | 0 | 0 | 0 | NA | 0 | 0 |
|  | 491 | M | 2 | 0 | + | 0 | 0 | 0 | 0 | 0 | 0 | 0 |
|  | 481 | F | 3 | 0 | + | 0 | 0 | 0 | 0 | NA | 0 | 0 |
|  | 490 | M | 3 | 0 | + | 0 | 0 | 0 | 0 | 0 | 0 | 0 |
|  | 484 | F | 5 | 0 | + | + | 0 | 0 | 0 | NA | 0 | 0 |
|  | 493 | M | 5 | 0 | + | 0 | 0 | 0 | 0 | 0 | 0 | 0 |
|  | 477 | F | 7 | 0 | 0 | 0 | 0 | 0 | 0 | NA | 0 | 0 |
|  | 485 | M | 7 | + | 0 | 0 | 0 | 0 | 0 | 0 | 0 | 0 |
|  | 476 | F | 9 | 0 | 0 | 0 | 0 | 0 | 0 | NA | 0 | + |
|  | 486 | M | 9 | + | 0 | 0 | 0 | 0 | 0 | 0 | 0 | + |

<sup>1</sup> Days PE - Days post exposure; + - SUDV-related macroscopic observation(s) noted; 0 - No SUDV-related macroscopic observations noted; NA - Not applicable; F – Female; M – Male; SE - Scheduled Euthanasia; SP-TK - Survival and Physiology Time Kinetics.

The earliest SUDV-related macroscopic observation was dark discoloration of the inguinal and axillary lymph nodes, noted on Day 2 PE. Enlarged axillary lymph nodes and friable dark red spleen were observed by Day 5. Cutaneous rash, discoloration of the mucosa in segments of the gastrointestinal tract, and discoloration at the exposure site were first noted on Day 7. Other less frequently noted macroscopic observations were found on or after Day 8, including pale liver, discoloration of testes, and red discoloration of the urinary bladder mucosa. Discoloration, scabbing and firmness at the exposure site, and enlarged, tan discolored axillary lymph nodes were observed in the sole surviving SUDV-exposed animal (Animal 487).

Animals exposed to SUDV presented with some common macroscopic findings (percentages based on SUDV-exposed animals only): 50% (n = 9) had axillary lymph node abnormalities (discoloration and/or enlargement and/or firmness); 44% (n = 8) had inguinal lymph node abnormalities (discoloration and/or enlargement); 39% (n = 7) had a skin rash; and 33% (n = 6) had red or black mucosa of the rectum. Some findings occurred in fewer than 5 animals: 17% (n = 3) had spleen abnormalities (discoloration and/or enlargement and/or friability); 17% (n = 3) had liver abnormalities (pale discoloration and/or enlargement); 11% (n = 2) exhibited red mucosa in the urinary bladder; and 11% (n = 2) exhibited firm and/or brown discoloration at the exposure site. Two males exhibited pink or red discoloration of the testes.

Other findings were less common at necropsy and were only observed in single animals. Animal 482 (unscheduled euthanasia on Day 8 PE with clinical score of 46) exhibited red discoloration of the dorsal nose. Animal 488 (unscheduled euthanasia on Day 9 with clinical score of 28) exhibited nasal hemorrhage and perianal fecal staining. Gastrointestinal findings were observed in Animal 492 (unscheduled euthanasia on Day 8 with clinical score of 24) and animal 494 (unscheduled euthanasia on Day 13 with clinical score of 20): dark green duodenum as noted in animal 492 and dark green or black contents in the stomach, jejunum, ileum, colon, and cecum were noted in animal 494. Animal 484 (scheduled euthanasia on Day 5 with clinical score of 3) exhibited diffuse red discoloration of the uterine endometrium.

#### Microscopic Observations

No SUDV-related microscopic changes were observed in the mock-exposed animals. In SUDV-exposed animals, the earliest SUDV-related findings were sinus histiocytosis and sinus erythrocytosis noted on Day 2 PE in axillary and inguinal lymph nodes.

SUDV-related findings at the Day 5 PE necropsy were noted in in lymph nodes, spleen, liver, adrenal gland, and at the exposure site (Figure 8 through Figure 10). Lymph node findings consisted of cortical lymphoid depletion, sinus histiocytosis, acute inflammation, and vasculitis with hemorrhage and fibrin deposition. Lymphoid depletion and fibrin were noted in the spleen. Findings within the liver consisted of hepatocellular necrosis and foci of inflammation. Single cell necrosis was noted in the cortex of adrenal glands. Inflammation and hemorrhage were noted in the skin and subjacent skeletal muscle at the exposure site.

**Figure 8.**
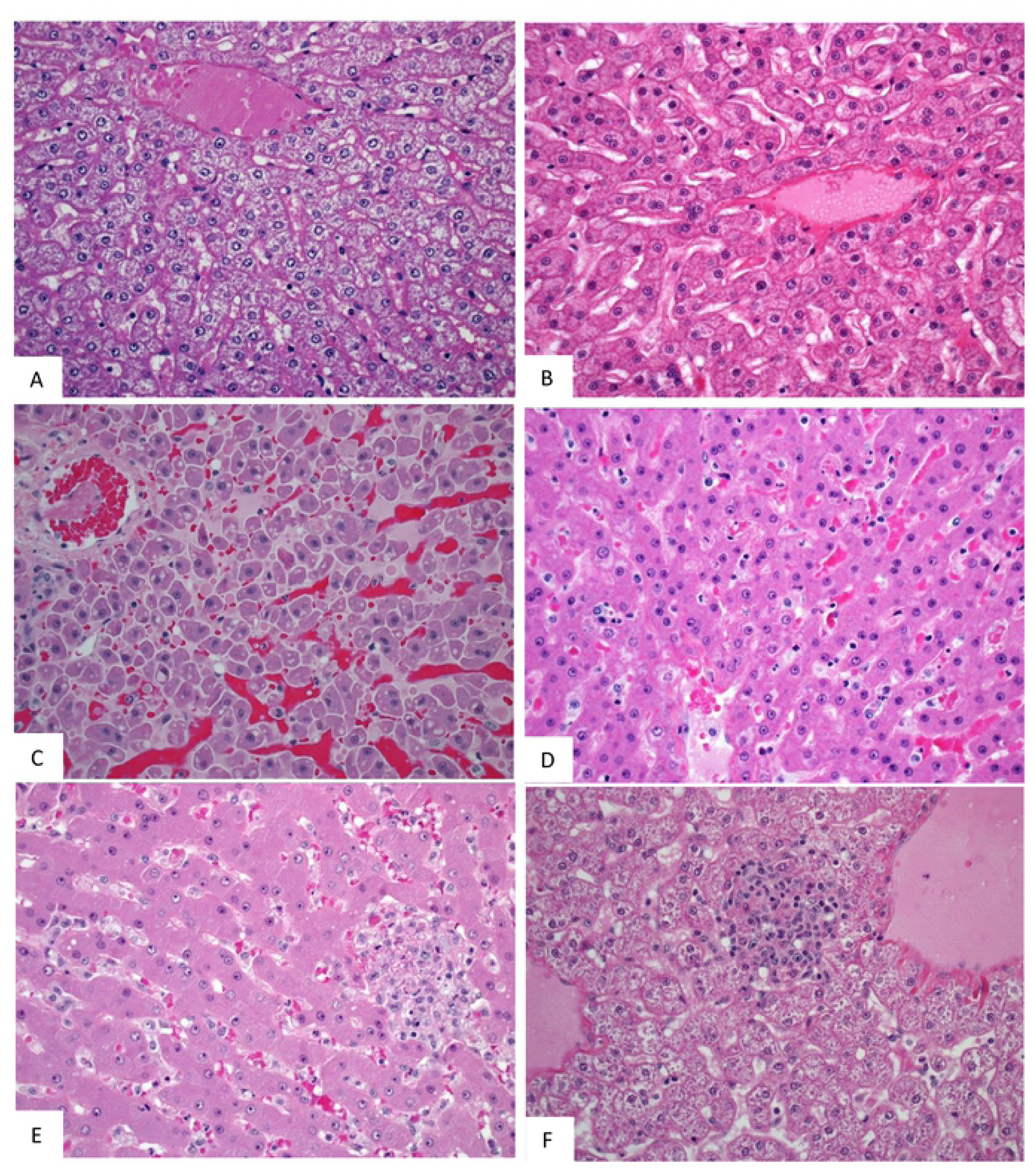
Representative images of microscopic findings: Axillary lymph node. Microscopic findings at necropsy in cynomolgus macaques intramuscularly exposed to SUDV Gulu. Axillary lymph node. (A) Day 21 control, No. 478. Essentially normal tissue. 20x; (A) Day 2 PE scheduled euthanasia, No. 491. There are increased histiocytes within subcapsular and medullary sinuses. 20x; (C) Day 5 PE scheduled euthanasia, No. 493. There is marked loss of lymphocytes with fibrin, hemorrhage, acute inflammation, edema, and increased histiocytes. 20x; (D) Day 7 PE scheduled euthanasia, No. 477. There is marked loss of lymphocytes with necrosis, hemorrhage, inflammation, fibrin, and increased histiocytes. 20x; (E) Day 9 PE scheduled necropsy, No. 476. There is lymphoid depletion with indistinct follicles, lymphocytolysis, and increased histiocytes. 20x; (F) Day 21 PE SUDV-exposed survivor, No. 487. Follicles are large with prominent germinal centers (follicular hyperplasia). Other findings include sinus histiocytosis, capsular fibrosis, and mononuclear cell inflammation. 10x.

**Figure 9.**
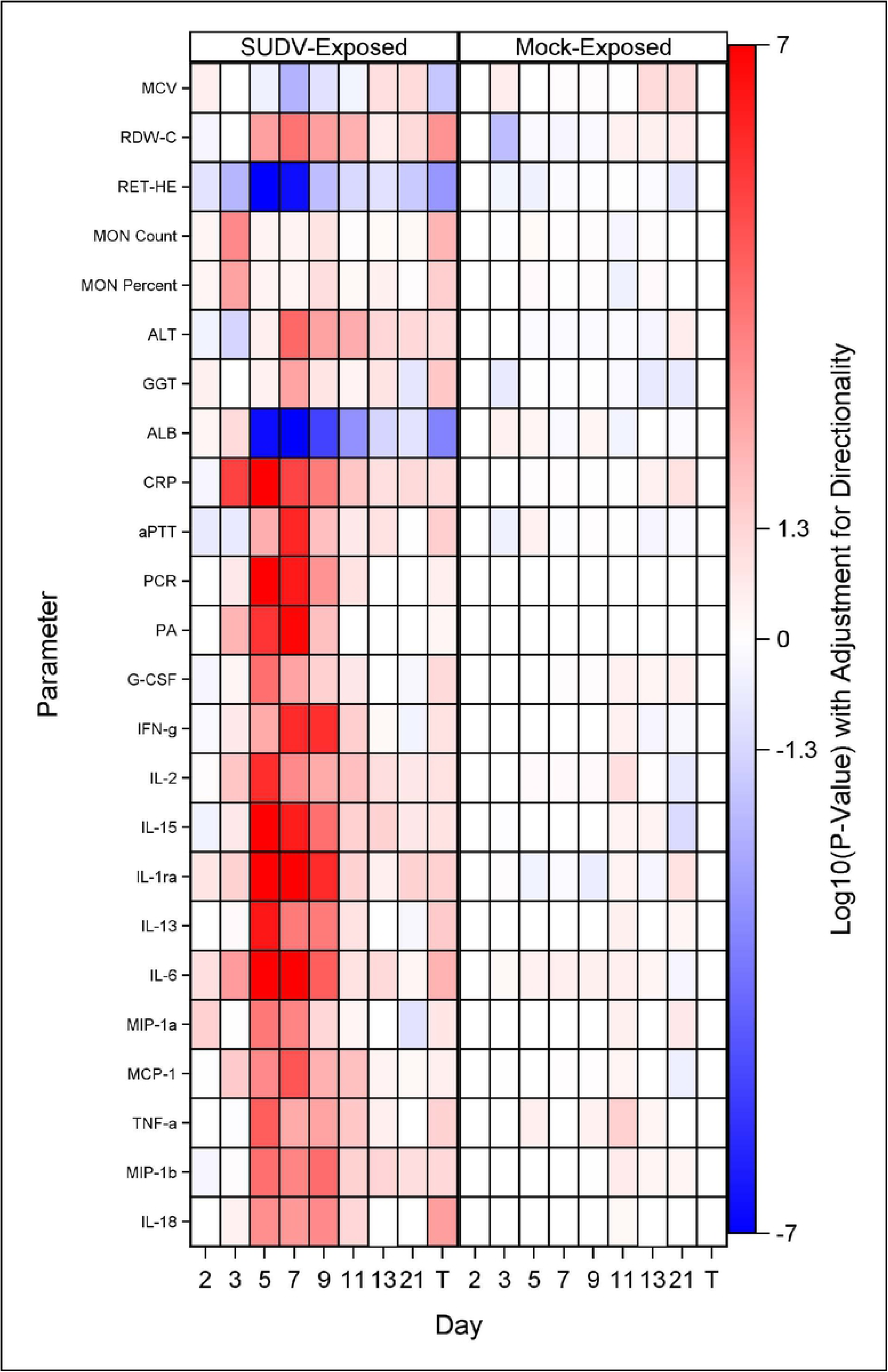
Representative images of microscopic findings: Spleen. Microscopic findings at necropsy in cynomolgus macaques intramuscularly exposed to SUDV Gulu. Spleen. (A) Day 21 control, No. 478. Essentially normal tissue. 4x; (B) Day 2 PE scheduled necropsy, No. 491. Follicles contain increased hyaline material, consistent with spontaneous change. 20x. (C) Day 5 PE scheduled euthanasia, No. 493. There is moderate lymphoid depletion with deposition of fibrin within the perifollicular marginal sinus. 20x; (D) Day 7 PE scheduled necropsy, No. 477. There is moderate lymphoid depletion with diffuse fibrin deposition. 20x; (E) Day 9 PE scheduled necropsy, No. 476. There is lymphoid depletion with lymphocytolysis and marginal sinus fibrin deposition that extends into the red pulp. 20x; (F) Day 21 PE SUDV-exposed survivor, No. 487. Follicles are enlarged with prominent germinal centers (follicular hyperplasia). 10x.

**Figure 10.**
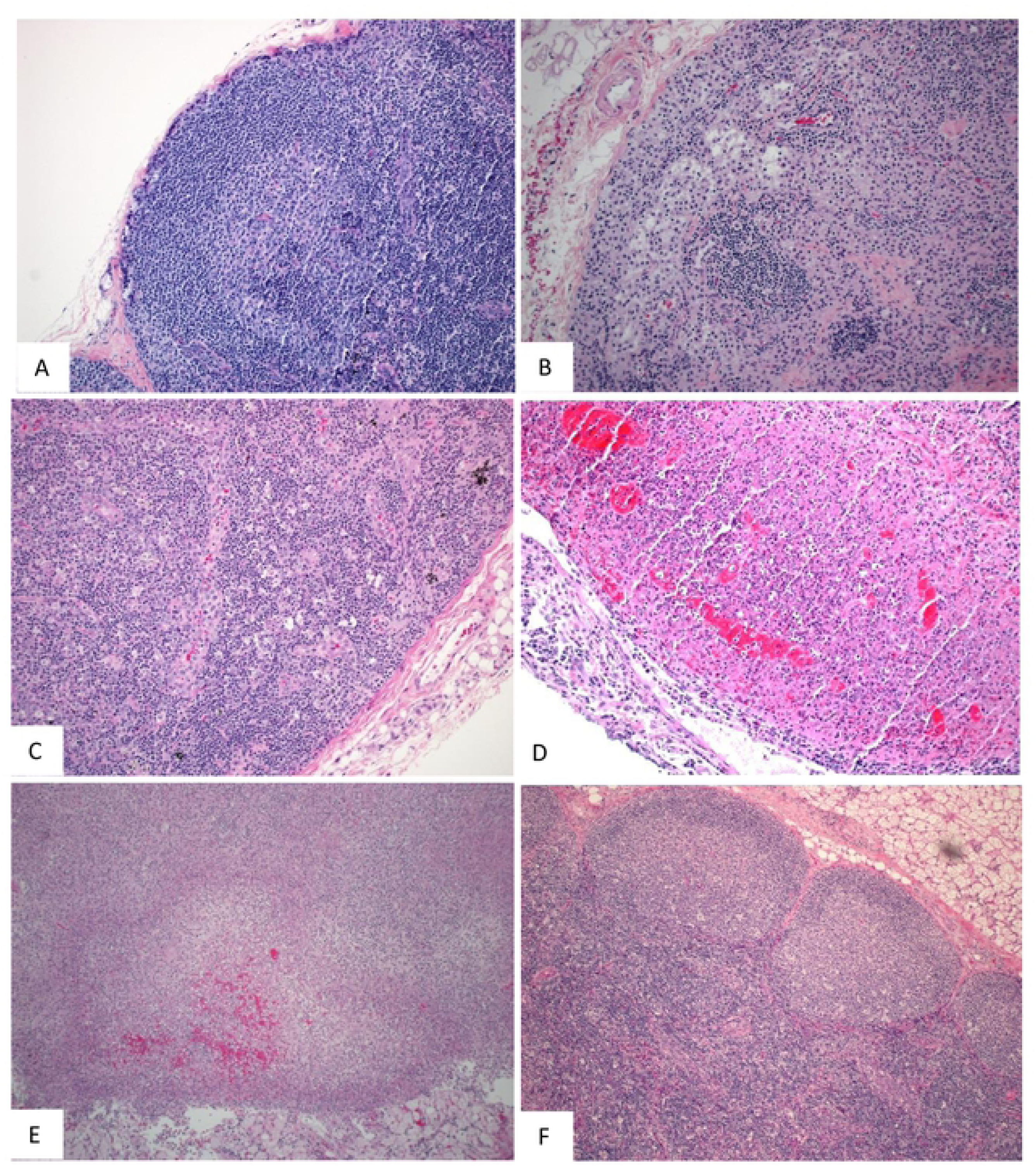
Representative images of microscopic findings: Liver. Microscopic findings at necropsy in cynomolgus macaques intramuscularly exposed to SUDV Gulu. Liver. (A) Day 21 control, No. 478. Essentially normal tissue. 40x; (B) Day 2 PE scheduled euthanasia, No. 491. Essentially normal tissue. 40x; (C) Day 5 PE scheduled euthanasia, No. 493. There is single cell hepatocellular degeneration, necrosis and vacuolation. 40x; (D) Day 7 PE, No. 477. There is rare necrosis of individual hepatocytes with inflammation and sinusoidal fibrin. 40x. (E) Day 9 PE scheduled euthanasia, No. 476. There is single cell hepatocellular necrosis with inflammation and fibrin. 40x; (F) Day 21 PE SUDV survivor, No. 487. There is multifocal mononuclear cell inflammation. 40x.

Microscopic findings noted at Day 5 PE persisted in the remaining SUDV-exposed monkeys euthanized or found dead from Day 7 to Day 13. Findings commonly noted in acutely fatal experimental filoviral infection in macaques were fully developed by Day 7. Systemic fibrin deposition, thrombosis, hemorrhage, inflammation, necrosis, lymphoid depletion, and lymphocytolysis were common microscopic features in monkeys from Day 7 to Day 13 PE. Fibrin thrombi within the microvasculature of duodenal Brunner’s glands were noted in most monkeys from Day 7 to Day 13 PE. Hemorrhage within segments of the gastrointestinal tract were additional common findings.

Microscopic findings in the single surviving SUDV-exposed monkey (Animal 487) consisted of inflammation, necrosis, fibrin, hemorrhage, and ulceration with serocellular crust at the exposure site. Minimal inflammation was noted in the liver. Lymphoid follicular hyperplasia was noted within the axillary lymph node and spleen, consistent with immune response to antigenic stimulation.

### Onset of Abnormality

There were few indications of SUDV infection until Day 5 post-exposure, when most animals had detectable infectious virus or viral RNA copies in serum. By Day 7, when animals began to succumb due to moribundity, there was universal evidence for infection from: virological data (plaque assay, qRT-PCR, sGP); gross abnormalities observed in multiple organs consistent with systemic infection; microscopic findings including inflammation, necrosis, fibrin, thrombosis, and hemorrhage noted in multiple organs; and other biomarkers commonly correlated with SUDV-induced disease including increased body temperature, elevated GGT, ALT, ALP, BA, BUN, CRP, and decreased ALB values. Statistical analysis (log rank tests) indicated that for time from exposure to onset of abnormality, there were statistically significant differences (at the 0.05 level) between SUDV-exposed animals and mock-exposed animals for 26 parameters (Figure 11). The earliest parameters for significant time to onset in SUDV- exposed animals were increased CRP and IL-6 on Day 3, followed by detection of infectious virus in the serum; increased IL-1ra, IFN- γ, and IL-15; and increased body temperature via telemetry on Day 4. By Day 5, numerous parameters were abnormal including clinical chemistry (ALT, GGT, ALB), hematology (MCV, RDW-C, RET-He, MON Count, and MON Percent), coagulation (aPTT), viral RNA (via qRT-PCR), sGP, and various cytokines/chemokines.

**Figure 11.**
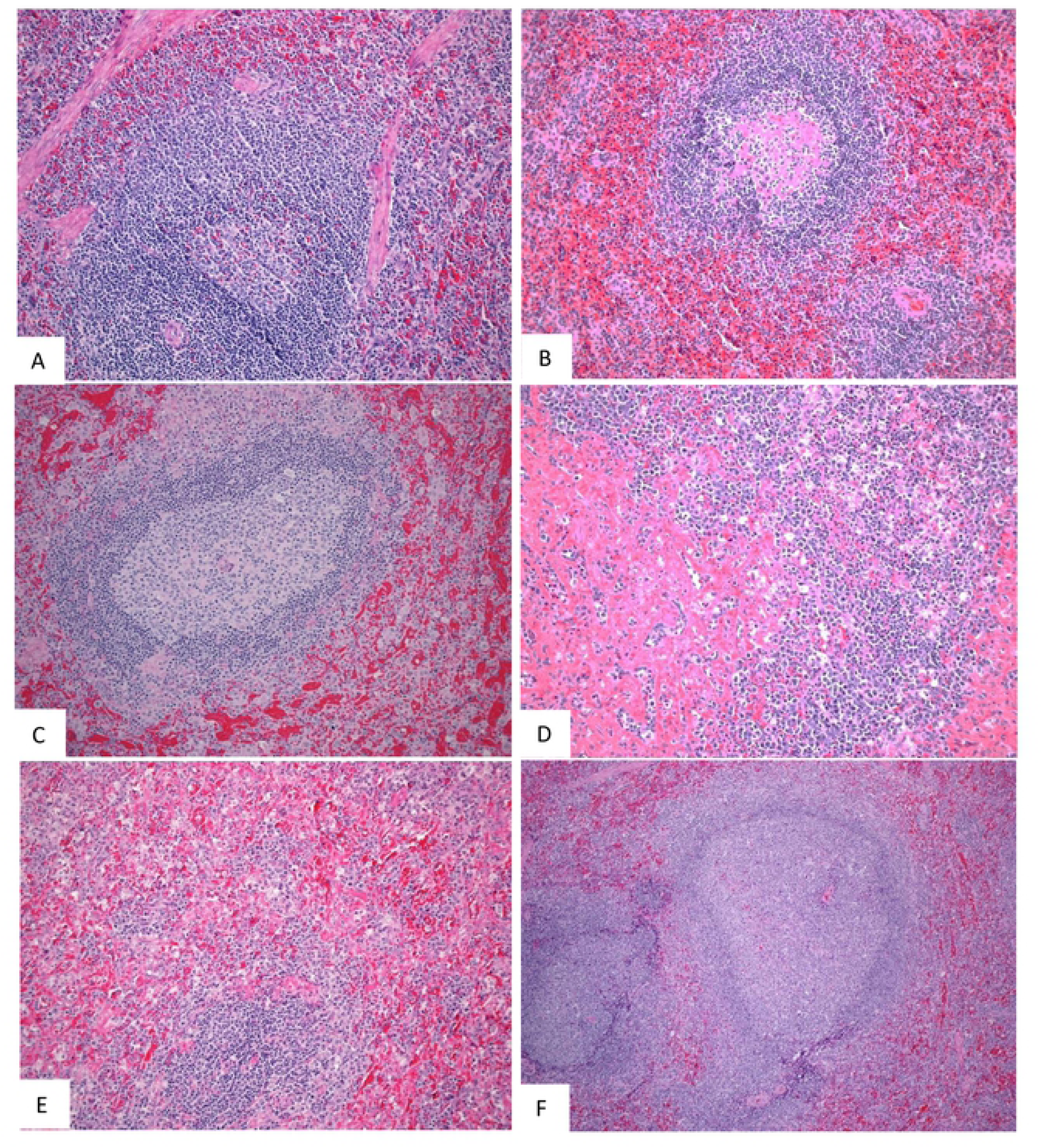
Onset of Abnormality Heat Map. T represents Terminal (data collected from animals that met euthanasia criteria were combined and reported as a single terminal time point).

Many of these parameters are consistent with the rhesus macaque model of EBOV exposure [18, 30]. In a similarly designed study targeted at characterizing EBOV exposure via the intramuscular route in the rhesus macaque, disease progression was marginally quicker for EBOV exposed animals than what was observed herein for SUDV exposed animals. Median time to death was one day quicker (Day 8 versus Day 9), clinical chemistry abnormalities and evidence of coagulopathy were more frequent one day earlier in EBOV exposed animals, body temperatures peaked earlier, and sGP was found in the serum of all animals earlier; serum viral load was similar between the two models Table 8 [18].

**Table 8.**
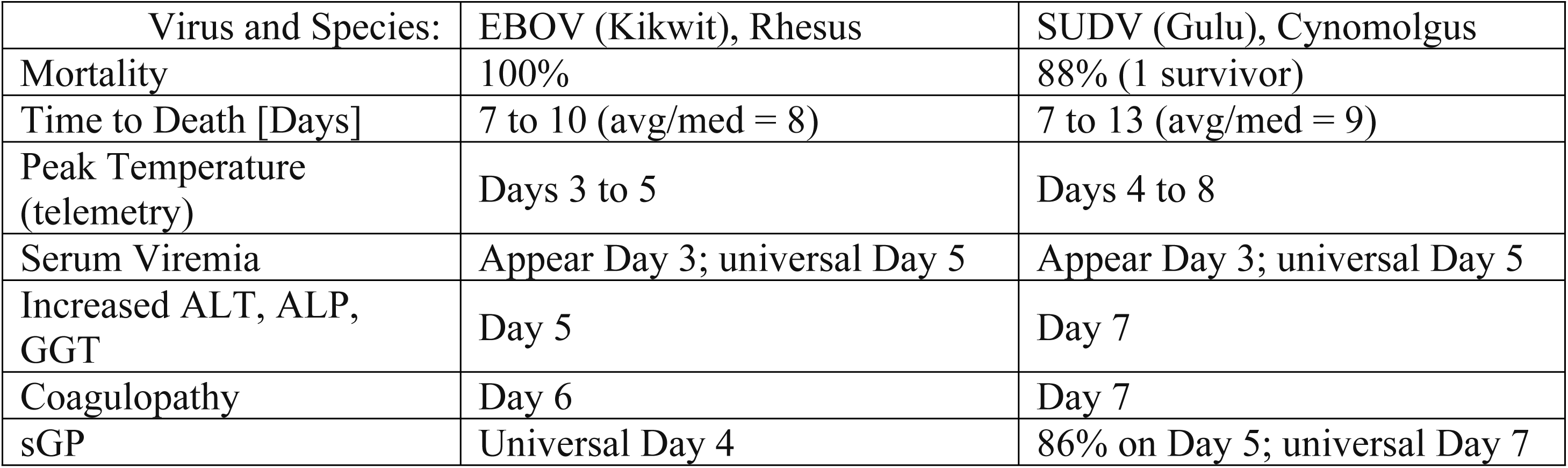
Comparison of Disease in Macaques Exposed via IM route to EBOV or SUDV.

| Virus and Species: | EBOV (Kikwit), Rhesus | SUDV (Gulu), Cynomolgus |
| --- | --- | --- |
| Mortality | 100% | 88% (1 survivor) |
| Time to Death [Days] | 7 to 10 (avg/med = 8) | 7 to 13 (avg/med = 9) |
| Peak Temperature (telemetry) | Days 3 to 5 | Days 4 to 8 |
| Serum Viremia | Appear Day 3; universal Day 5 | Appear Day 3; universal Day 5 |
| Increased ALT, ALP, GGT | Day 5 | Day 7 |
| Coagulopathy | Day 6 | Day 7 |
| sGP | Universal Day 4 | 86% on Day 5; universal Day 7 |

## Discussion

The primary objective of this study was to characterize the disease course in cynomolgus macaques exposed via the intramuscular route to a target dose of 1000 PFU SUDV Gulu variant to determine if infection in this species is an appropriate model for the evaluation of filovirus countermeasures under the FDA Animal Rule. The Gulu variant utilized in this study has been identified as a suitable isolate for exposure material in nonclinical testing and evaluation under the Animal Rule [24].

Two mock-exposed animals were housed at ABSL-4 and euthanized on Day 21 post exposure. During most observations, these animals exhibited no clinical signs, and infrequent low clinical scores were assigned for reduced feed consumption or stool abnormalities. Throughout the study, these animals exhibited few notable abnormalities in body temperature, body weight, or clinical pathology parameters. Mock-exposed animals were normal at gross examination and no significant microscopic lesions were noted.

For virus-exposed animals, numerous virologic and clinical parameters were observed during SDV progression. SUDV (infectious virus and viral RNA) were first detected in serum on Day 3 and most animals were positive for these parameters by Day 5. SUDV sGP is produced during infection and has a putative role in pathogenicity [31], and detection of this biomarker in serum was consistent with the presence of virus. Clinical signs of disease were minimal until Day 5 when reduced responsiveness and increased rectal temperatures in individual animals were observed. As the time post-exposure progressed, reduced responsiveness, food and water intake, and stool output were commonly observed, along with the presence of petechia and increased rectal temperatures. Symptoms consistently observed shortly before moribund euthanasia included severely reduced responsiveness, hypothermia, bleeding at a site other than the collection site, and nasal discharge.

Indicators of multi-organ system involvement of SVD were observed around the same time that the clinical symptoms were observed. Clinical chemistry parameters indicative of marked liver damage, such as ALP, ALT, GGT, BA, and TBIL are frequently associated with SVD in humans and NHPs [12, 25, 26, 32] and were abnormal (significant increase or decrease from baseline values) beginning around Day 5 to 7. Additionally, up to 3-fold increases in BUN and decreased ALB [25] was observed consistently in the exposed animal group beginning around Day 5 and extending throughout the remainder of the disease process. Indicators of coagulation abnormalities were also commonly observed, with platelet counts decreased on Day 3 and throughout the remainder of the study period and increased clotting times (aPTT and PT) by Day 5. CRP, a marker of inflammation, was significantly increased by Day 3 and maintained high levels throughout the disease period.

The immunological response observed in the SUDV-exposed animals was consistent with observations in human filovirus infections: high levels of pro-inflammatory cytokines (e.g. IL- 1β, IL-4, IL-1RA, IL-6, IL-8, IL-15 and IL-16) and chemokines (e.g. MIP-1α, MIP-1β, MCP-1) observed in human infections [27, 32] were observed during acute infection and peak disease around Days 5 to 9 post-exposure. In addition, high levels of sCD40L have been detected in survivors leading to the suggestion that sCD40L could be a novel biomarker to predict clinical outcome [28]. NHPs in this study exhibited increased levels of most cytokines and chemokines analyzed, similar to what has been reported in humans; decreased levels of sCD40L were also observed, which is expected in non-survivors based on the human data.

The earliest macroscopic findings were darkly discolored axillary and inguinal lymph nodes, which correlated with sinus erythrocytosis, consistent with lymphoid drainage of a site of hemorrhage. By Day 5, enlarged lymph nodes were observed and this finding correlated with a depletion of lymphocytes in the spleen and lymph nodes. The appearance of hepatocellular and adrenocortical cell necrosis, inflammation at the exposure site, and fibrin deposition were also observed. Enlarged spleen was observed on Day 7, which correlated with fibrin deposition. Day 8 observations included pale liver, which correlated with hepatocellular necrosis. Common macroscopic findings observed in SUDV-exposed animals at moribund necropsy included axillary and inguinal lymph node abnormalities (discolored and/or enlarged and/or firm) and red or black mucosa of the rectum; these findings correlated with high titers of virus. As the infection spreads and becomes more systemic, evidence for the damage associated with the disease was detected in other organs, especially in the spleen and liver where there is major blood flow. Finally, gross pathology abnormalities became more common in multiple organs for the animals euthanized at the latter stages of the disease.

Animal 487 that survived infection had detectable levels of infectious SUDV in serum on Days 5, 7, and 9, and sGP in serum was detected on Days 7, 9, 11, and 13. The animal exhibited some clinical signs of infection beginning on Day 5 and throughout the timeframe where other animals succumbed, including reduced responsiveness, increased temperature, and reduced intake of fluid and/or food. Numerous abnormalities in hematology and clinical chemistry parameters were observed for animal 487; however, increased GGT, BA, BUN, and CHOL were not observed in this animal but were consistently observed in animals that succumbed. At necropsy (Day 21), the surviving animal exhibited macroscopic and microscopic findings at the exposure site consistent with SUDV infection. Enlarged and tan discolored axillary lymph nodes observed macroscopically correlated with lymphoid follicular hyperplasia, consistent with immune response to antigenic stimulation; follicular hyperplasia was additionally noted in the spleen. Hepatic inflammation was also noted microscopically. The proinflammatory cytokine IL- 17 peaked in this animal on Day 7 post exposure. The role of IL-17 and Th17 cells in filovirus infections remains an area of investigation [26], but it is possible the spike in IL-17 had a protective effect in this animal.

The scheduled euthanasia in this study provided the opportunity to study the earliest stages of the disease in order to determine if the measured biomarkers may be correlated with clinical signs and the pathological spread of the disease. The emphasis on the early stages of the disease also provides an opportunity to identify triggers that are appropriate for intervention in a therapeutic model.

This study was limited by the small number of animals used, a common problem for animal modeling studies performed in high containment laboratories, due to resource limitations; however, the results show the model reproducibly develops disease in most exposed animals. In addition, future studies should utilize a longer timeframe – if possible, given the constraints of performing long term studies at BSL-4 – especially in light of what is being learned about EBOV long term sequelae and persistence.

The data in this study support the observation that IM exposure of cynomolgus macaques to 1000 PFU SUDV Gulu results in a rapid systemic disease similar to the infection in humans. The local nature of the initial disease means there are no unambiguous universal indicators of infection until Day 5 to 7, at which time a number of biomarkers were available as evidence for infection including RNA copy number by qRT-PCR, liver enzyme elevation, increased clotting time, and changes in cytokines/chemokines.

## Materials and Methods

### Ethics Statement

Animal research was conducted under an Institutional Animal Care and Use Committee (IACUC)-approved protocol (IACUC number 1634MF) in compliance with the Animal Welfare Act, Public Health Service (PHS) policy, and other federal statutes and regulations relating to animals and experiments involving animals. Texas Biomedical Research Institute (Texas Biomed) is accredited by AAALAC International. Euthanasia criteria were developed to minimize undue pain and distress and animals were euthanized with an overdose of sodium pentobarbital after study veterinarian approval.

### Critical Biological Materials

Sudan virus (SUDV) Gulu variant was used for animal exposures and was supplied by Texas Biomed. A second cell-culture passage (P2) of *Sudan ebolavirus* Gulu was obtained from Dr. Tom Ksiazek (at National Institute of Allergy and Infectious Diseases (NIAID’s) World Reference Center for Emerging Viruses and Arboviruses (WRCEVA) at the University of Texas Medical Branch (UTMB) Health Galveston National Laboratory) in 2012 and passaged for a third time in Vero E6 cells [12, 25]. Sterile PBS (Gibco) was used to mock-expose control animals.

### Test System Experimental History

Twenty (20) Chinese origin cynomolgus macaques (*Macaca fascicularis*), 10 male and 10 female, were used in this study. On the day of exposure, animals were 4.7 to 5.7 years of age and weighed 2.78 to 8.07 kg. Animals were acquired from Envigo (previously Covance; Alice, TX) 61 days prior to exposure. Prior to study enrollment, NHPs were verified to be: experimentally naïve; seronegative for Simian Immunodeficiency Virus (SIV), Simian T-Lymphotropic Virus-1 (STLV-1), Simian Varicella Virus (SVV) and *Macacine herpesvirus* 1 (Herpes B virus); PCR negative for Simian Retrovirus (SRV1 and SRV2); negative for *Trypanosoma cruzi* (PCR and serology); free from active infections with *Salmonella* and *Shigella*; negative for tuberculosis; antibody-negative for Ebola Reston nucleoprotein (screened by Virus Reference Laboratory, San Antonio, TX, USA); and antibody-negative for Ebola virus, Sudan virus, and Marburg virus glycoprotein (screened at Texas Biomed). Animals were implanted with M00 telemeter implants (DSI) at Texas Biomed by a Southwest National Primate Research Center (SNPRC) veterinarian and surgical veterinary technician staff. The telemeters were implanted intra-abdominally in the subperitoneal space, according to an IACUC approved protocol. Preoperatively, animals received Buprenorphine SR and meloxicam via subcutaneous route. Immediately prior to the surgery, animals were sedated with Telazol via the IM route. During surgery, animals were anesthetized with inhalational isoflurane (0.5%-2.0%). Postoperatively, animals received Clavamox and meloxicam.

### Animal Care

Animals were housed individually in stainless steel cages with wire mesh bottoms and sides. Excreta pans under the cages, cage flooring, and room floors were cleaned daily. Animals were fed commercially available primate diet from Purina Mills (Diet 5048) one to two times daily, and at least five times per week were provided additional edible enrichment. Water from the Institutional Watering System was available *ad libitum*. Structural perches and toys were provided as inanimate enrichment. Environmental and photoperiod conditions were: temperature range of 74°F ± 10°F, humidity range of approximately 30 to 70%, and light cycle of approximately 12 hours on/12 hours off. Light cycle conditions were interrupted for extended observations beginning on Day 5 post exposure and continuing through Day 13 post exposure.

Animals were evaluated by a study veterinarian to confirm health prior to transfer to the ABSL-4. Animals were observed by veterinary technician staff at least twice daily at least 6 hours ± 2 hours apart for morbidity and mortality. Clinical observations involved evaluating each animal for thirteen different parameters and assigning a numerical score to each parameter [18, 33]. Briefly, the following parameters were assessed: feed, enrichment, and fluid consumption, with reduced consumption warranting a score of 1 or 2; stool output, with abnormal output warranting a score of 1 or 2; hair coat appearance (rough hair coat was assigned a score of 1); presence of nasal discharge (score of 1); presence of bleeding, with bleeding assigned a score of 1 or 2 based on the source. Other parameters were weighted more heavily: respiration was observed to determine if breathing was normal (score of 0), labored (score of 8), or agonal (score of 15); and responsiveness was observed to determine if the animal had diminished activity (score of 1), reduced response to external stimuli (score of 2), moderate to dramatically reduced response to stimuli (score of 8), or was severely/completely unresponsive (score of 15). On days when animals were sedated, they were also assessed for changes in rectal body temperature, decreased body weight, and the presence of petechia. Scores were then added up to achieve a total clinical score, which was reported to the study veterinarian when above a 3. A clinical score of 4 to 7 in any animal resulted in all animals being observed at least three times per day and a clinical score greater than 7 in any animal resulted in all animals being observed at least four times a day.

Animals were euthanized when the total clinical score reached 15, or if they scored 8 for responsiveness and also exhibited either a greater than 5 degree temperature change or increases above a predetermined range in two or more of certain clinical chemistry parameters [18, 33]. Any animals found moribund were euthanized with the approval of the responsible veterinarian.

### Exposure Agent Preparation, Administration, and Verification

Animals were acclimated to the ABSL-4 for 24 days prior to exposure. Animals were exposed to a target dose of 1,000 PFU SUDV Gulu diluted in sterile PBS. Mock-exposed animals received undiluted sterile PBS. A total of 0.5 mL of exposure material was administered intramuscularly (IM) to each animal in the right deltoid muscle of the arm. Prior to virus or PBS injection: NHPs were sedated via IM injection with Telazol (Zoetis), body weight and rectal temperature were recorded, and a blood sample was collected. Following exposure, each NHP was taken back to its home cage and observed until it had recovered from sedation.

Following preparation of the exposure material, aliquots were removed for determination of viral titer by plaque assay. The titer for mock-infection material was below the limit of detection (as expected for control material). The titer for SUDV exposure material was 1,630 PFU per 0.5 mL. After the last animal was exposed, the viral titer of the exposure material remaining after injection was also confirmed by plaque assay. The post-exposure titer for mock-infection material was below the limit of detection (as expected for the control material). The post-exposure titer for SUDV exposure material was 1,810 PFU per 0.5 mL.

### Blinding and Randomization

Veterinary staff (technicians and veterinarians) were blinded to group assignment (i.e. SUDV exposed versus mock-exposed with PBS, for animals not in the scheduled euthanasia cohort) until finalization of post-in life analysis. In vitro staff, performing viral load via RT-qPCR and cytokine analysis were blinded to group assignment (i.e. SUDV exposed versus mock-exposed with PBS, for animals not in the scheduled euthanasia cohort); staff were not blinded regarding the time point (day post exposure) of each sample when they performed the assays and analysis. The individuals performing necropsy were blinded to group assignment (i.e. SUDV exposed versus mock-exposed with PBS, for animals not in the scheduled euthanasia cohort). The board-certified veterinary pathologist was blinded to the animal group assignments during microscopic evaluation of H&E stained slides, and unblinded for preparation of the pathology report and microscopic evaluation of immunohistochemistry slides. Animals were randomly assigned to one of four groups (two groups of eight animals and two groups of two animals) in order to ensure that group size did not indicate which group was the mock-exposed control group.

### Blood Collection and Analysis

Blood samples were collected via femoral or saphenous venipuncture on Study Days -24, 0, 3, 5, 7, 9, 11, 13, and 21 and prior to euthanasia. Whole blood was collected from a vein into plastic serum separator tube for coagulation analysis, and subsequently processed to obtain serum. Whole blood was also collected into tubes containing EDTA for hematology analysis and clinical chemistry, and subsequently processed to obtain plasma. Complete Blood Counts were performed using a ProCyte Dx Hematology Analyzer. Evaluation of the coagulation parameters activated partial thromboplastin time (aPTT) and prothrombin time (PT) was performed using an IDEXX Coag Dx Analyzer. Clinical chemistry analysis was performed using Mammalian Liver Profile and Abaxis Comprehensive Diagnostic Panel rotors on VetScan analyzers. The following parameters were analyzed: alkaline phosphatase (ALP), alanine aminotransferase (ALT), gamma-glutamyl transferase (GGT), bile acids (BA), total bilirubin (TBIL), albumin (ALB), blood urea nitrogen (BUN), and cholesterol (CHOL), amylase (AMY), calcium (CA2+), creatinine (CRE), globulin (GLOB), glucose (GLU), potassium (K+), sodium (NA+), phosphorus (PHOS), and total protein (TP). In addition, serum was analyzed using a Piccolo® BioChemistry Panel Plus rotor to determine C-Reactive Protein (CRP) levels.

### Virus Quantification via Plaque Assay

Viremia was determined at the completion of the in-life portion of the study on serum collected from Study Day 0 up to the day of euthanasia. Viral load was determined on samples taken from 15 tissues collected at necropsy. For liver and spleen, two samples (excised from different sites of the organ) were analyzed via plaque assay to assess variability. If considerable differences in titer were found between the two samples, a third sample was assayed for viral load by plaque assay. Finally, back titrations were performed on the exposure material. Virus quantification was performed by plaque assay, using neutral red and crystal violet agarose overlay [34].

### Virus Quantification via Quantitative Reverse Transcription Polymerase Chain Reaction

RNA copy number was determined by qRT-PCR targeting a region of the SUDV glycoprotein. The primer and probe information for this qRT-PCR assay are as follows: Forward Primer: 5’ CCA CTC TCA CCA CCC CAG AA 3’; Reverse Primer: 5’ ACC CGT GGC TTT GGT GTT AG 3’; Probe: 6-FAM 5’ TTG GGC TTC GAA AAC GCA GCA GAA 3’. Serum samples were inactivated using RNAbee Reagent (Tel-Test, Friendswood, TX), following manufacturer instructions. Tissue samples were inactivated using Trizol Reagent (Invitrogen).

### Cytokine analysis

The MILLIPLEX MAP Non-Human Primate Cytokine Magnetic Bead Panel was used, along with Luminex xMAP detection, to examine the cytokine profiles in serum. Analytes assayed included the following: Granulocyte-colony stimulating factor (G-CSF), granulocyte-macrophage colony-stimulating factor (GM-CSF), interferon gamma (IFN-γ), interleukin (IL)- 1ra, IL-1β, IL-2, IL-4, IL-5, IL-6, IL-8, IL-10, IL-12/23 (p40), IL-13, IL-15, IL-17, IL-18, monocyte chemotactic protein-1 (MCP-1), macrophage inflammatory protein (MIP)-1α, MIP-1β, soluble CD40 ligand (sCD40L), transforming growth factor alpha (TGF-α), tumor necrosis factor alpha (TNF-α), and vascular endothelial growth factor (VEGF).

### Soluble Glycoprotein (sGP) Analysis

Serum was analyzed for soluble GP using an ELISA (IBT Bioservices, SUDV Soluble GP (sGP) ELISA Kit, catalog number 0102-001). It has been reported that the manufacturer recommended standard curve (range of 0.105 ng/mL to 1,000 ng/mL) begins to plateau at the higher concentrations, resulting in less reliable quantification [18]. Thus, the manufacturer recommended method was modified to use a truncated standard curve with a range of 1.56 to 100 ng/mL. Samples were tested at dilutions of 1:50, 1:200, 1:500, or 1:25,000 depending on the time point tested. The initial 1:50 dilution was prepared in manufacturer recommended buffer and any subsequent dilutions were prepared in recommended buffer supplemented with 2% cynomolgus macaque serum (BioIVT, Westbury, NY, USA) in order to maintain a consistent serum concentration across all dilutions. A quality control sample of sGP prepared in cynomolgus macaque serum at a known concentration was also included to evaluate data quality and consistency. Given the standard curve range of 1.56 to 100 ng/mL and the minimum required dilution of 1:50, the range of quantification with this assay is 78 to 5,000 ng/mL. However, the process for diluting samples eliminates the upper limit of quantification.

### Body Temperature, Body Weight, and Activity Data Collection

Rectal body temperatures and body weights were measured for each animal at least once prior to transfer to ABSL-4, when they were sedated for blood collection, on the day of euthanasia and during a cage change on Days -11 or -12. Prior to collection, animals were sedated via IM injection of Telazol.

Body temperature data were also collected from M00 telemeter implants. Upon study completion, data collected from the implants were uploaded to an online enterprise content management platform called Box for access by DSI personnel. After exposure, hourly averages were compared to the baseline established before exposure (72 hours, three days prior to SUDV exposure). Data were only processed if the signals were of sufficient quality to be analyzed. Data were omitted in the case of signal drop out and/or non-physiological values. Parameters evaluated were: Temperature, Temperature-mean, Activity, Mean activity.

### Necropsy and Pathology Analysis

All animals euthanized were subjected to a complete necropsy no later than 12 hours after euthanasia. Animal carcasses held longer than 1 hour after euthanasia and prior to necropsy were refrigerated. The necropsy included examination and recording of findings of the external surfaces of the body, all orifices, and the cranial, thoracic and abdominal cavities and their contents. Macroscopic necropsy observations were recorded using consistent descriptive terminology to document location(s), size, shape, color, consistency, and number of lesions. Samples of tissues were aseptically removed and divided for viral load determination and stored frozen, or fixed in 10% neutral-buffered formalin.

Tissues were fixed by immersion in 10% neutral-buffered formalin for a minimum of 14 days, then trimmed, routinely processed, and embedded in paraffin. Embedded tissues were sectioned at 5 um thick and histology slides were deparaffinized, stained with hematoxylin and eosin (H & E), coverslipped, and labelled. Slides were evaluated by a board-certified veterinary pathologist.

### Data Analysis and Statistics

Preliminary analysis was performed to assess the model assumption of normality, to identify potential outliers, and to determine whether there are significant differences between the groups at baseline. ANOVA models were fitted to the change from baseline data to determine if there were significant changes from the pre-exposure baseline within a group, or differences between the groups. Statistical analyses were conducted using SAS® (version 9.4; SAS, Cary, NC, USA) on the 64-bit platform. Results are reported at the 0.05 level of significance.

### Quality System

This study adhered to a thorough Study Protocol, Standard Operating Procedures (SOPs), generally recognized good documentation practices, and a quality agreement that assigned roles and responsibilities of study staff and was consistent with GLP principals. Further quality measures were as previously described [18] and were consistent with FDA Animal Rule Guidelines for adequate and well controlled studies to ensure data quality and integrity.

## Acknowledgments

The authors would like to acknowledge Wendy Melendez, Shawn Santos, and Leah McDonald for their assistance in reviewing data. Special thanks to the veterinary staff for their assistance with the *in vivo* exposure study: Laura Rumpf, Matthew Stautzenberger, Tony Bowers, Alex Limon, Sadye Steele, and Drs. Edward Dick and Olga Gonzalez. Dr. Thomas Ksiazek at the University of Texas Medical Branch kindly provided the P2 virus.

## Funding

This project has been funded in whole or in part with federal funds from the Department of Health and Human Services; Office of the Assistant Secretary for Preparedness and Response; Biomedical Advanced Research and Development Authority, under Contract No. HHSO100201700011I. The views expressed here are those of the authors and do not necessarily represent the views or official position of the Funding agencies. This work was conducted in facilities constructed with support from the Research Facilities Improvement Program (grant number C06 RR012087) from the NCRR. This investigation used resources that were supported by the Southwest National Primate Research Center grant P51 OD011133 from the Office of Research Infrastructure Programs, National Institutes of Health.

## Author contributions

Y.G. and K.J.A contributed equally and designed and conducted experiments and wrote the paper. H.S. and M.G. conducted and designed experiments and analyzed data. M.M. analyzed macroscopic and microscopic pathology data, and assisted in writing. E.A.C. and J.W.D analyzed clinical pathology data. A.T., B.K., G.R., and P.E. conducted experiments and analyzed data. N.N. and Y.C. performed statistical analysis. C.M.C., G.T.M., and D.C.S. designed experiments, analyzed data, and assisted in writing. A.G. and R.C. designed experiments and analyzed data.

## Competing interests/Conflicts of Interest

The authors declare not conflicts of interest..

## Notes

### Competing Interest Statement

The authors have declared no competing interest.

